# *Cis*-regulatory Element Hijacking by Structural Variants Overshadows Higher-Order Topological Changes in Prostate Cancer

**DOI:** 10.1101/2021.01.05.425333

**Authors:** James R. Hawley, Stanley Zhou, Christopher Arlidge, Giacomo Grillo, Ken Kron, Rupert Hugh-White, Theodorus van der Kwast, Michael Fraser, Paul C. Boutros, Robert G. Bristow, Mathieu Lupien

## Abstract

Prostate cancer is a heterogeneous disease whose progression is linked to genome instability. However the impact of this instability on the three-dimensional chromatin organization and how this drives progression is unclear. Using primary benign and tumour tissue, we find a high concordance in the higher-order three-dimensional genome organization across normal and prostate cancer cells. This concordance argues for constraints to the topology of prostate tumour genomes. Nonetheless, we identify changes to focal chromatin interactions and show how structural variants can induce these changes to guide *cis*-regulatory element hijacking. Such events result in opposing differential expression on genes found at antipodes of rearrangements. Collectively, our results argue that *cis*-regulatory element hijacking from structural variant-induced altered focal chromatin interactions overshadows higher-order topological changes in the development of primary prostate cancer.

## Introduction

The human genome is organized into hubs of chromatin interactions within the nucleus, setting its three-dimensional topology ^1^. Two classes of higher-order topology, topologically associating domains (TADs) and compartments, define clusters of contacts between DNA elements that are linearly distant from each other, such as *cis*-regulatory elements (CREs) and their target gene promoters ^2, 3^. Insulating these hubs to prevent ectopic interactions are TAD boundaries, maintained by CCCTC-binding Factor (CTCF) and the cohesin complex ^4^. Disruption of TAD boundaries through genetic or epigenetic variants can activate oncogenes, as observed in medulloblastoma^5^, acute myeloid leukemia^6^, gliomas ^7^, and salivary gland acinic cell carcinoma ^8^. However, recent studies depleting CTCF or the cohesin complex produced little effect on gene expression despite global changes to the three-dimensional chromatin organization ^9–11^. In contrast, CRE hijacking caused by genetic alterations can result in large changes to gene expression, despite having little impact on the higher-order chromatin organization ^5, 12^. These contrasting observations raise questions about the interplay between components of the genetic architecture, namely, how genetic alterations, chromatin states, and the three-dimensional genome cooperate to misregulate genes in disease. Understanding the roles that chromatin organization and *cis-*regulatory interactions play in gene regulation is crucial for understanding how their disruption can promote oncogenesis.

The roles of noncoding mutations targeting CREs in cancer are becoming increasingly clear ^12–14^. Mutations to the *TERT* promoter, for example, lead to its over-expression and telomere elongation in multiple cancer types ^15–17^. Similarly, mutations targeting CREs of the *ESR1* and *FOXA1* oncogenes in breast and prostate cancers, respectively, lead to their sustained over-expression ^18–20^, which is associated with resistance to hormonal therapies ^21–24^. Point mutations have the potential to alter three-dimensional chromatin organization, albeit indirectly, by modifying transcription factor or CTCF binding sites ^25, 26^. Structural variants (SVs), on the other hand, are large rearrangements of chromatin that can directly impact its structure ^27, 28^. This can establish novel CRE interactions from separate TADs or chromosomes, as has been observed in leukemia ^29^ and multiple developmental diseases ^30, 31^. But how prevalent and to what extent these rearrangements affect the surrounding chromatin remains largely unstudied in primary tumours ^14, 28, 32^. Hence, to understand gene misregulation in cancer, it is critical to understand how SVs impact three-dimensional chromatin organization and CRE interactions in primary tumours.

SVs play an important role in prostate cancer (PCa), both for oncogenesis and progression. An estimated 97% of primary tumours contain SVs ^14, 33^, and translocations and duplications of CREs for oncogenes such as *AR* ^34^, *ERG* ^35^, *FOXA1* ^20, 36^ and *MYC* ^20^ are highly recurrent. While coding mutations of *FOXA1* are found in ∼10% of metastatic castration-resistant PCa patients, SVs that target *FOXA1* CREs are found in over 25% of metastatic prostate tumours ^20^. In addition to oncogenic activation, SVs in prostate tumours disrupt and inactivate key tumour suppressor genes including *PTEN, BRCA2, CDK12,* and *TP53* ^36, 37^. Furthermore, over 90% of prostate tumours contain complex SVs, including chromothripsis and chromoplexy events ^38^, making it a prime model to study the effects of SVs. However, despite large-scale tumour sequencing efforts, investigating the impact of SVs on three-dimensional prostate genome remains difficult, owing to constraints from chromatin conformation capture (i.e. Hi-C) assays. In this work, we build on recent technological advances in Hi-C protocols to investigate the three-dimensional chromatin organization of the prostate from primary benign and tumour tissues. Using patient-matched whole genome sequencing (WGS), RNA sequencing (RNA-seq), and chromatin immunoprecipitation (ChIP-seq) data, we show that SVs in PCa repeatedly hijacking CREs to disrupt the expression of multiple genes with minimal impact to higher-order three-dimensional chromatin organization.

## Results

### Three-dimensional Chromatin organization is stable over oncogenesis

Chromatin conformation capture technologies enable the measurement of three-dimensional chromatin organization. These assays, however, are often limited to cell lines, animal models and liquid tumours due to the amount of input required^39^. Here, we optimized and conducted low-input Hi-C ^40^ on 10 μm thick cryosections from 12 primary prostate tumours and 5 primary benign prostate sections (see Methods, Figure 1a, Supplementary Figure 1a). The 12 tumours were selected from the Canadian Prostate Cancer Genome Network (CPC-GENE) cohort previously assessed for whole-genome sequencing ^33^, RNA-seq ^41^ and H3K27ac ChIP-seq ^42, 43^ (Supplementary Table 1). All 12 of these PCa patients previously underwent radical prostatectomies and 6 of our 12 samples (50%) harbour the *TMPRSS2-ERG* genetic fusion (T2E) found in approximately half of the primary PCa patients ^33^. The total percent of genome altered ranges from 0.99%-18.78% (Supplementary Table 1) ^33^. The 12 tumour samples were histopathologically assessed to have ≥70% cellularity while the cellularity was ≥ 60% for our group of 5 normal prostate samples. Upon Hi-C sequencing, we reached an average of 9.90 x 10^8^ read pairs per sample (range 5.84 x 10^8^ -1.49 x 10^9^ read pairs) with minimal duplication rates (range 10.6% - 20.8%) (Supplementary Table 2). Pre-processing resulted in an average of 6.23x10^8^ (96.13%) valid read pairs per sample (range 3.95x10^8^ - 9.01x10^8^, or 82.42 - 99.22%; Supplementary Table 2). Hence, we produced a high depth, high quality Hi-C library on 17 primary prostate tissue slices.

**Figure 1.**
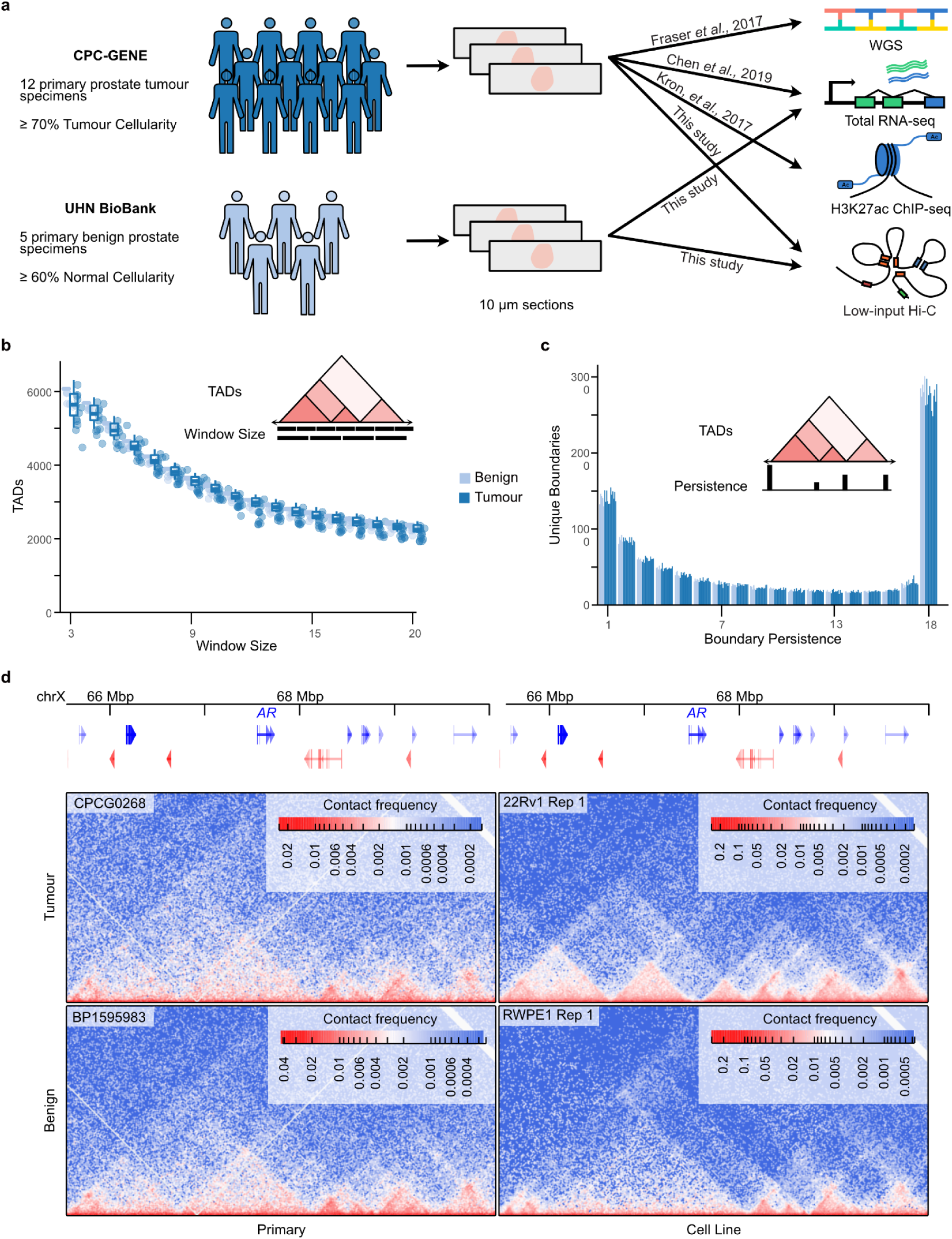
Topologically associated domains are stable over prostate oncogenesis. **a**. The sample collection and data usage of primary prostate samples in this study. 10 µm sections from 6 tumours previously identified as T2E+ and 6 T2E- were used for Hi-C sequencing. 5 additional 10 µm sections were collected from benign prostate specimens in the UHN BioBank. **b-c.** A comparison of the number of TADs detected at multiple window sizes (**b**) and boundary persistence (**c**) in each patient sample, with inset schematics. **d.** Contact matrices around the *AR* gene demonstrate a difference in chromatin organization between primary samples and cell lines. Hi-C data for 22Rv1 and RWPE1 cell lines obtained from^44^.

To characterize the higher-order organization of the primary prostate genome, we first identified TADs. Across the 17 primary tissue samples, we observed an average of 2,305 TADs with a median size of 560 kbp (Supplementary Tables 3-4). However, when considering all hierarchical levels of TAD organization, we did not observe significant differences in the number of TADs identified across length scales (Figure 1b), nor in the persistence of their boundaries (Figure 1c). This suggests few, if any, differences in three-dimensional chromatin organization at the TAD level between benign and tumour tissue. However, we observed differences in organization around essential genes for PCa between primary tissue and previously profiled cell lines. For example, chromatin around the *AR* gene that was previously found enriched in the 22Rv1 compared to RWPE1 prostate cell lines ^44^ were not recapitulated in either benign or tumour primary samples (Figure 1d, Supplementary Figure 1d). Moreover, when compared to other Hi-C datasets, the primary prostate samples clustered separately from cell lines (Supplementary Figure 1b), despite similar enrichment of CTCF binding sites near TAD boundaries (Supplementary Figure 1c). These results suggest that TADs are constrained over oncogenesis and that cell line models may not harbour disease-relevant three-dimensional chromatin organization.

We next investigated compartmentalization changes, the second class of higher-order three-dimensional chromatin organization. Recurrent changes to segments nearly the size of chromosome arms showed differential compartmentalization in multiple tumour samples compared to benign samples, such as compartment B-to-A transitions on 19q and A-to-B transitions on chromosome Y (Supplementary Figure 2a-c). Only two genes on chromosome 19 were differentially expressed between the 8 tumours with benign-like compartmentalization and the other 4 (Supplementary Figure 2d). Similarly, no genes on chromosome Y were differentially expressed between the 4 tumours with benign-like compartmentalization and the remaining samples (Supplementary Figure 2e). Both arms on chromosome 3 show differential mean compartmentalization, but this appears to be driven by one tumour sample and one benign sample for each arm and is not recurrent (Supplementary Figure 2f). Collectively, these results suggest that phenotypic differences between benign and tumour tissues do not stem from differences in higher-order three-dimensional chromatin organization alone.

### Focal chromatin interactions shifting over oncogenesis

Changes to focal chromatin interactions have been observed in the absence of higher-order chromatin changes ^45, 46^, and we hypothesized that this may be the case in PCa. We detected chromatin interactions, identifying a median of 4,395 interactions per sample (range 1,286 - 6,993; Supplementary Figure 3a, Supplementary Table 5). Among these detected interactions, we identified known contacts in PCa such as those between two distal CREs on chromosome 14 and the *FOXA1* promoter ^19^ (Supplementary Figure 3b), and CREs upstream of *MYC* on chromosome 8 that are frequently duplicated in metastatic disease ^36^ (Supplementary Table 5). 16,474 unique chromatin interactions were identified in at least one sample (Figure 2a), reaching an estimated ∼80% saturation of detection (Supplementary Figure 3c). Restricting our analysis to the 8,486 interactions present in at least two samples (51.5% of all interactions) yielded 1,405 tumour- and 273 benign-specific interactions, suggesting focal changes in three-dimensional chromatin organization occur over oncogenesis. Aggregate peak analysis revealed Hi-C contact enrichment at all detected interactions in all samples (Figure 2b-c), demonstrating that tumors- and benign-specific interactions are not binary. Rather, the contacts at “tumour-specific” loci are more enriched than those at “benign-specific” loci in tumour samples (Figure 2b). Similarly, the contacts at “benign-specific” loci are more enriched than those at “tumour-specific” loci in benign samples (Figure 2c). Together, these results suggest that more focal changes to chromatin interactions are present in prostate oncogenesis despite the stable higher-order organization.

**Figure 2.**
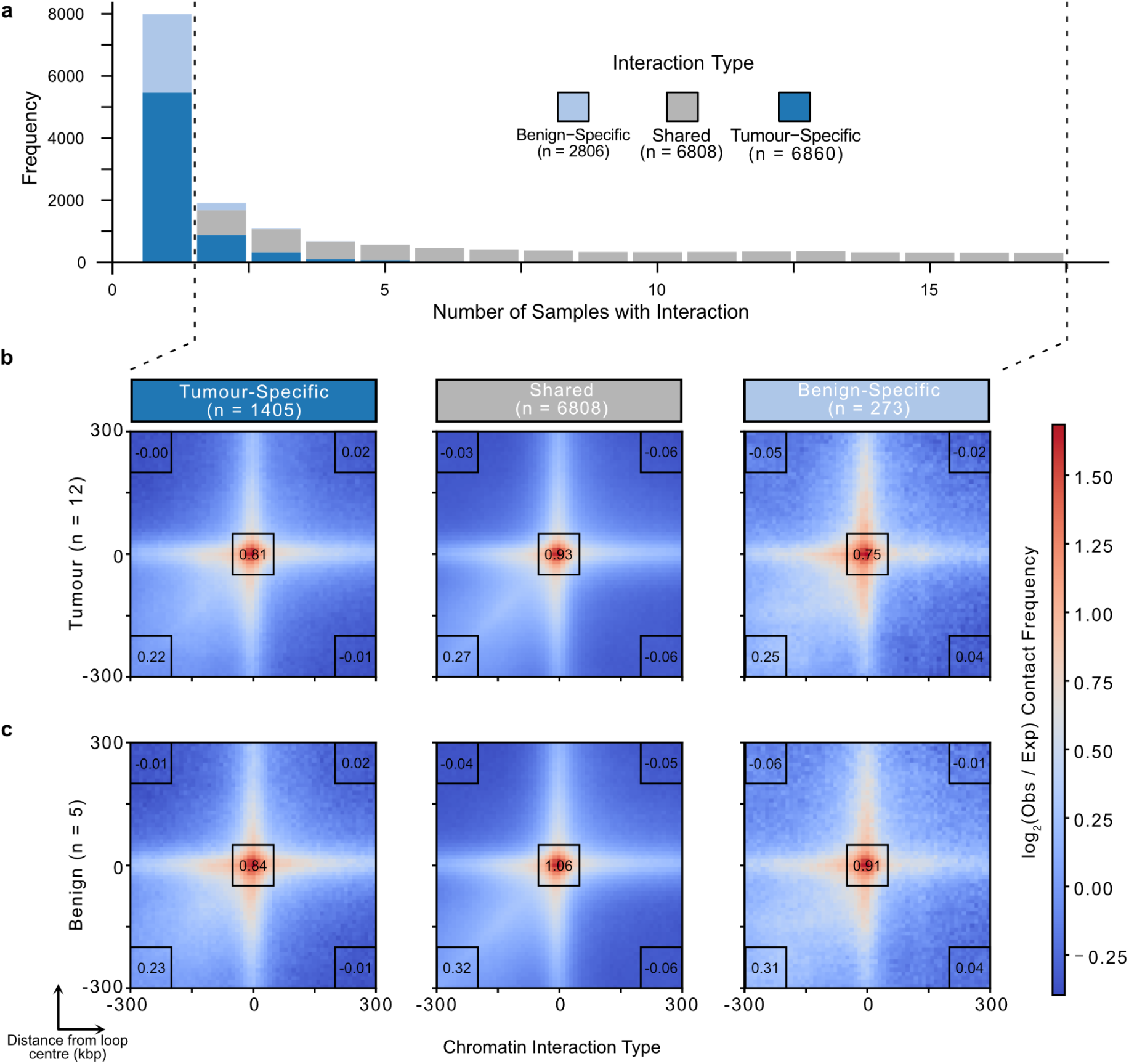
Focal chromatin interactions display subtle differences between benign and tumour tissue. **a.** Stacked bar plots of the number of samples chromatin interactions were identified in. **b-c.** Aggregate peak analysis of tumour (**b**) or benign (**c**) contacts in tumour-specific, benign-specific, and shared interactions identified in two or more samples. Regions plotted are +/- 300 kbp around the centre of each identified interaction. Inset numbers are the mean log2(obs/exp) contact frequencies within the 100 kbp x 100 kbp black boxes.

### Cataloguing structural variants from Hi-C data

In prostate tumours, SVs populate the genome to aid disease onset and progression ^33, 36^. Advances in computational methods now enable the identification of SVs from Hi-C datasets ^27, 47^. Applying an SV caller to our primary prostate tumour Hi-C dataset^27^, we detected a total of 317 unique breakpoints with a median of 15 unique breakpoints per tumour (range 3-95; Figure 3a; Supplementary Table 6). As an example, we found evidence of the *TMPRSS2-ERG* (T2E) genetic fusion spanning the 21q22.2-3 locus in 6/12 (50%) patients (CPCG0258, CPCG0324, CPCG0331, CPCG0336, CPCG0342, and CPCG0366) (Figure 3b), in accordance with previous whole-genome sequencing (WGS) findings ^33^. Combining unique breakpoint pairs into rearrangement events yielded 7.5 total events on average per patient (range 1 - 36, Supplementary Figure 4a-b). We also identified more inter-chromosomal breakpoint pairs with the Hi-C data in 11 of 12 tumours (Figure 3b), including a novel translocation event that encompasses the deleted region between *TMPRSS2* and *ERG* into chromosome 14. Few loci contained SV breakpoints recurrent between patients (Supplementary Figure 4c). These numbers are smaller than previously reported from matched WGS data ^33^; however, the median distance between breakpoints on the same chromosome was much larger at 31.6 Mbp for Hi-C-identified breakpoints, compared to 1.47 Mbp from WGS-identified breakpoints (Figure 3c). This is consistent with the inherent nature and resolution of the Hi-C method to detect larger, inter-chromosomal events ^27^. No SVs were detected in the 5 primary benign prostate tissue samples from Hi-C data. While this does not rule out the presence of small rearrangements undetectable by Hi-C limited by its resolution, the absence of large and inter-chromosomal SVs further supports a difference in genome stability between benign and tumour tissues ^33, 38, 43, 48^. Collectively, Hi-C defines a valid method to interrogate for the presence of SV in tumour samples, compatible with the detection of intra- and inter-chromosomal interactions otherwise missed in WGS analyses.

**Figure 3.**
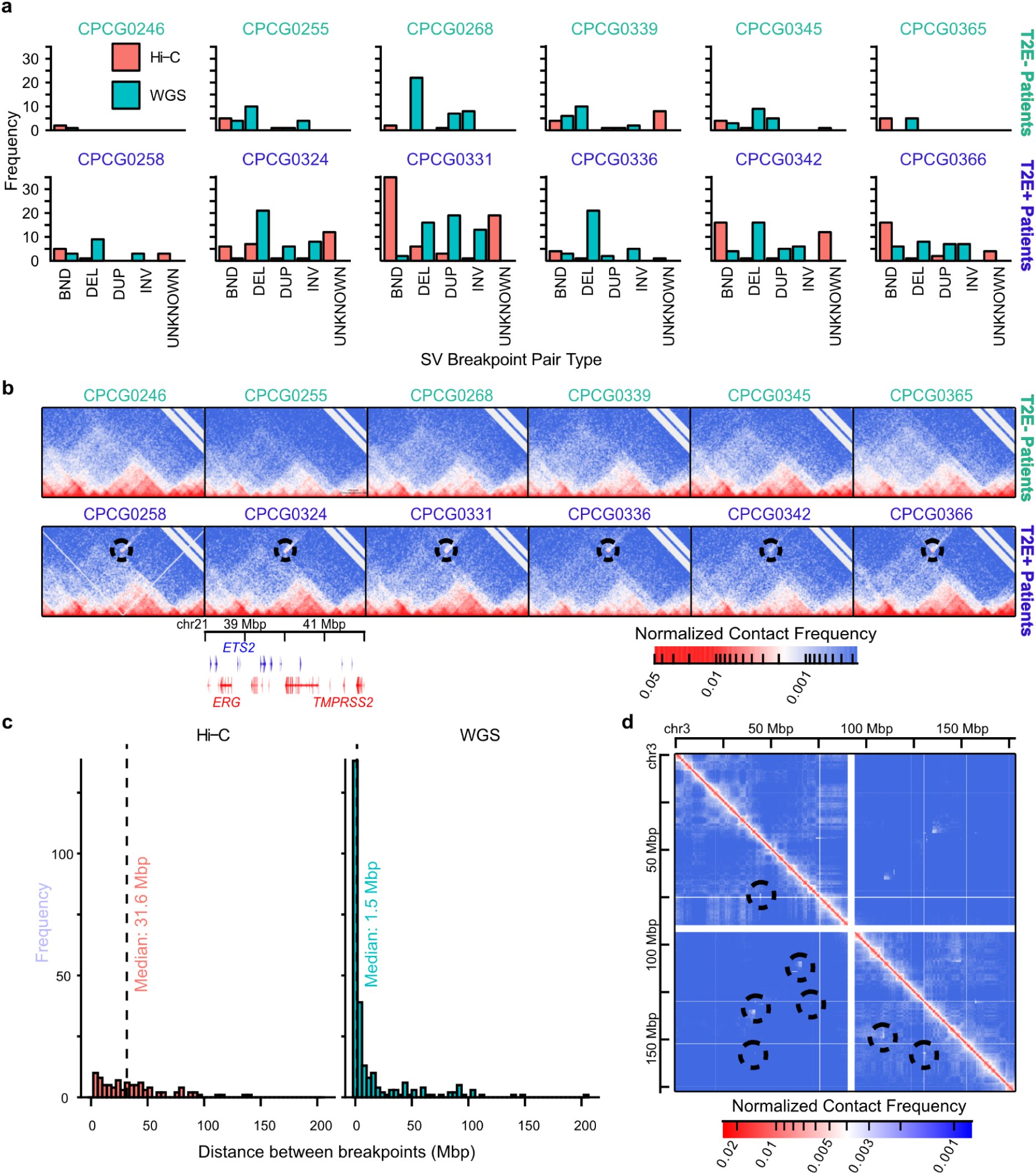
SVs are identified in primary tissue through chromatin conformation capture. **a.** Barplot of SV breakpoint pairs identified by Hi-C and WGS^33^ on matched samples. BND = inter-chromosomal translocation, DEL = deletion, DUP = duplication, INV = inversion, UNKNOWN = breakpoint pair of unknown type. **b.** Hi-C contact matrices of the chr21:37-42 Mbp locus harbouring the *TMPRSS2* and *ERG* genes. Circles indicate increased contact between *TMPRSS2* and *ERG* in the T2E+ tumours. **c.** Histogram showing the distance between breakpoints on the same chromosome detected by Hi-C (left) versus WGS^33^ (right). **d.** An example of a complex set of rearrangement spanning both arms of chromosome 3 in a patient.

Among SVs detected in primary prostate tumours, we identified both simple and complex chains of breakpoints. While simple SVs correspond to fusion between two distal DNA sequences, complex chains are evidence of chromothripsis and chromoplexy ^38^. These genomic aberrations affecting multiple regions of the genome are known to occur in both primary and metastatic PCa ^14, 33, 38^. The chains can be pictured as paths connecting breakpoints in the contact matrix (Supplementary Figure 4d). 8 of the 12 (66.7%) tumour samples contained these chains, including one patient (CPCG0331) harbouring 11 complex events and three patients (CPCG0246, CPCG0345, and CPCG0365) each harbouring a single complex event. We observed a median of 1 complex event per patient (range 0-11) consisting of a median of 3 breakpoints (range 3-7) spanning a median of 2 chromosomes per event (range 1-4, Supplementary Table 7, Supplementary Figure 4b). Patient CPCG0331 had 11 complex events, including a 6-breakpoint event spanning 3 chromosomes (Supplementary Figure 4b). A highly rearranged chromosome 3 was also found in the same patient (Figure 3d). The most common type of complex event involved 3 breakpoints and spanned 2 chromosomes, occurring 9 times across 5 of the 8 patients with complex events. In summary, using Hi-C, we detected both simple and complex SVs in primary prostate tumours not previously identified using WGS-based methods. We were able to identify known observations, such as a highly mutated region on chromosome 3 and subtype-specific differences in abundance, as well as find novel inter-chromosomal events not previously reported.

### SVs alter gene expression independently of intra-TAD contacts

Using combined WGS called SVs with those from Hi-C data, we next systematically examined the impact of SVs on TAD structure. This led us to look at the intra-TAD and inter-TAD interactions around each breakpoint. We observed that only 18 of the 260 (6.9%) TADs containing SV breakpoints were associated with decreased intra-TAD or increased inter-TAD interactions (Figure 4a). 12 of 18 (66.7%) occurrences were within T2E+ tumours. We found no evidence that simple versus complex SVs were a factor in determining whether a TAD was altered (Pearson’s chi-square test, X^2^ = 0.0166, p = 0.8974, df = 1). Similarly, the type of SV (a deletion, inversion, duplication, or translocation) was not predictive of whether the TAD would be altered (Pearson’s chi-square test, X^2^ = 4.7756, p = 0.3111, df = 4). Overall, we find that SVs are associated with higher-order topological changes in a small percentage of cases and that the presence of an SV breakpoint is not predictive alone of an altered TAD.

**Figure 4.**
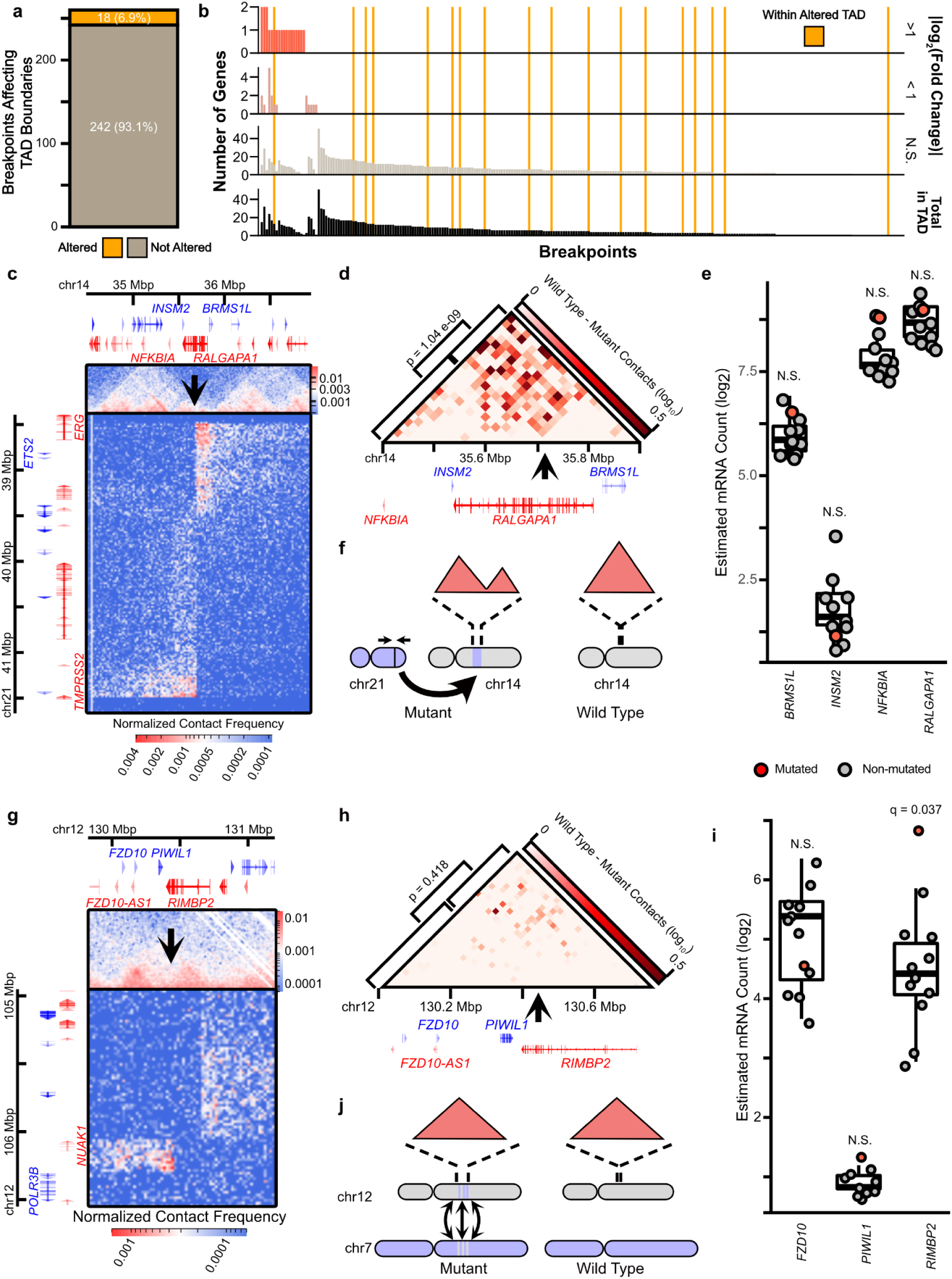
SVs can alter TADs or gene expression around breakpoints, but rarely alters both. **a.** A count of the number of SV breakpoints associated with altered TAD boundaries. **b.** Bar plot showing the number of genes differentially expressed around SV breakpoints. **c-f.** An example of an SV that alters TAD boundaries without significantly affecting gene expression of the nearby genes. **c.** The contact matrix showing a translocation of the *TMPRSS2-ERG* locus into chr14 in the *RALGAPA1* gene. **d.** The differential contact matrix between the tumour containing this translocation and another tumour without it to show the decreased contacts between sites upstream and downstream of the insertion site. **e.** Expression of the genes within the broken TAD show no significant changes to their expression. **f.** A schematic representation of the translocation. **g-j.** An example of an SV that does not alter TAD boundaries but does alter the expression of a nearby gene. **g.** The contact matrix showing a complex rearrangement around the *RIMBP2* gene. **h.** The differential contact matrix between the tumour containing this translocation and another tumour without it to show the decreased contacts between sites upstream and downstream of the insertion site. **i.** Expression of the genes within the broken TAD show no significant changes to their expression. **j.** A schematic representation of the translocation.

Despite the evidence that SVs rarely impact higher-order chromatin topology, we evaluated whether SVs affected the expression of genes within the TADs surrounding the breakpoint using patient-matched RNA-seq data ^41^. We found that 23 of 260 breakpoints (8.8%) are associated with significant changes to local gene expression (Figure 4b). Complex events can have opposite effects at each breakpoint. For example, while the T2E fusion across all tumours leads to over-expression of *ERG* and under-expression of *TMPRSS2* ^33, 42^, the deleted locus between these two genes was inserted into chromosome 14 as part of a complex translocation event in one patient (Figure 4c-f). This inserted fragment positions *ERG* towards the 5’ end of the *RALGAPA1* gene and *TMPRSS2* towards the 3’ end (Figure 4c) resulting in a significant drop in intra-TAD contacts at the *RALGAPA1* locus on chromosome 14 (two-sample unpaired t-test, t = 6.38, p=1.04e-9; Figure 4d). Despite the significant topological change on chromosome 14, no significant changes to expression was detectable across genes within the same TAD on chromosome 14 (Figure 4e). Conversely, TAD alterations are not required changes to gene expression. As part of a complex SV involving the *RIMBP2* gene (Figure 4g-j), both ends of the gene contain breakpoints (Figure 4g). This rearrangement is not associated with changes to intra-TAD contacts (two-sample unpaired t-test, t = 0.8101, p = 0.4183; Figure 4h). However, *RIMBP2* is over-expressed in this patient (Figure 4i). More generally, only a single breakpoint was observed with both TAD contact and gene expression changes, although we did not find evidence to suggest these are dependent events (Pearson’s chi-square test, X^2^ = 6.31e-3, p = 0.9367, df = 1). For TADs where at least one gene was differentially expressed, 19 (83%) of them had at least one gene with doubled or halved expression. Notably, we found that inter-chromosomal translocations are associated with altering the expression of genes nearby their breakpoints compared to intra-chromosomal breakpoints (Pearson’s chi-squared test, X^2^ = 7.0088, p = 0.00811, df = 1; Supplementary Figure 5). Taken together, these results suggest that while SVs can alter contacts within TADs, this is neither necessary nor sufficient to alter gene expression.

### SVs alter focal chromatin interactions to hijack CREs and alter antipode gene expression

Mutations in prostate cancer have previously been found to converge on active CREs ^49^. To assess if SVs function in a similar fashion, we investigated the convergence of SV breakpoints in active CREs. We find that SV breakpoints are enriched in the catalogue of CREs captured by H3K27ac ChIP-seq from our 12 primary prostate tumours compared to the rest of the genome (one-sided permutation z-test, z = 25.591, p = 0.0099, n = 100; Figure 5a-b). This is similar to the enrichment of point mutations in CREs active in prostate cancer ^49^, suggesting that SVs which alter gene expression may do so by recurrently targeting CREs. Since individual CREs can regulate multiple genes ^50^, we suspected that SVs that do alter gene expression may predominantly affect multiple genes at the same time, instead of single genes. In agreement, when considering all SVs associated with altered gene expression near a breakpoint we find 16 of the 22 (72.7%) SVs are associated with altered expression of multiple genes (Figure 5c-d). Notably, 15 of these 16 SVs (93.8%) are associated with both over- and under-expression of genes, instead of genes all being either over-expressed or under-expressed (Figure 5d). 12 of these 15 (80%) SVs are associated with expression changes at SV antipodes, opposite ends of a breakpoint pair (Supplementary Figure 6). The recurrent targeting of active CREs, combined with the opposite gene expression changes at SV antipodes, suggests that SVs may repeatedly alter expression by CRE hijacking.

**Figure 5.**
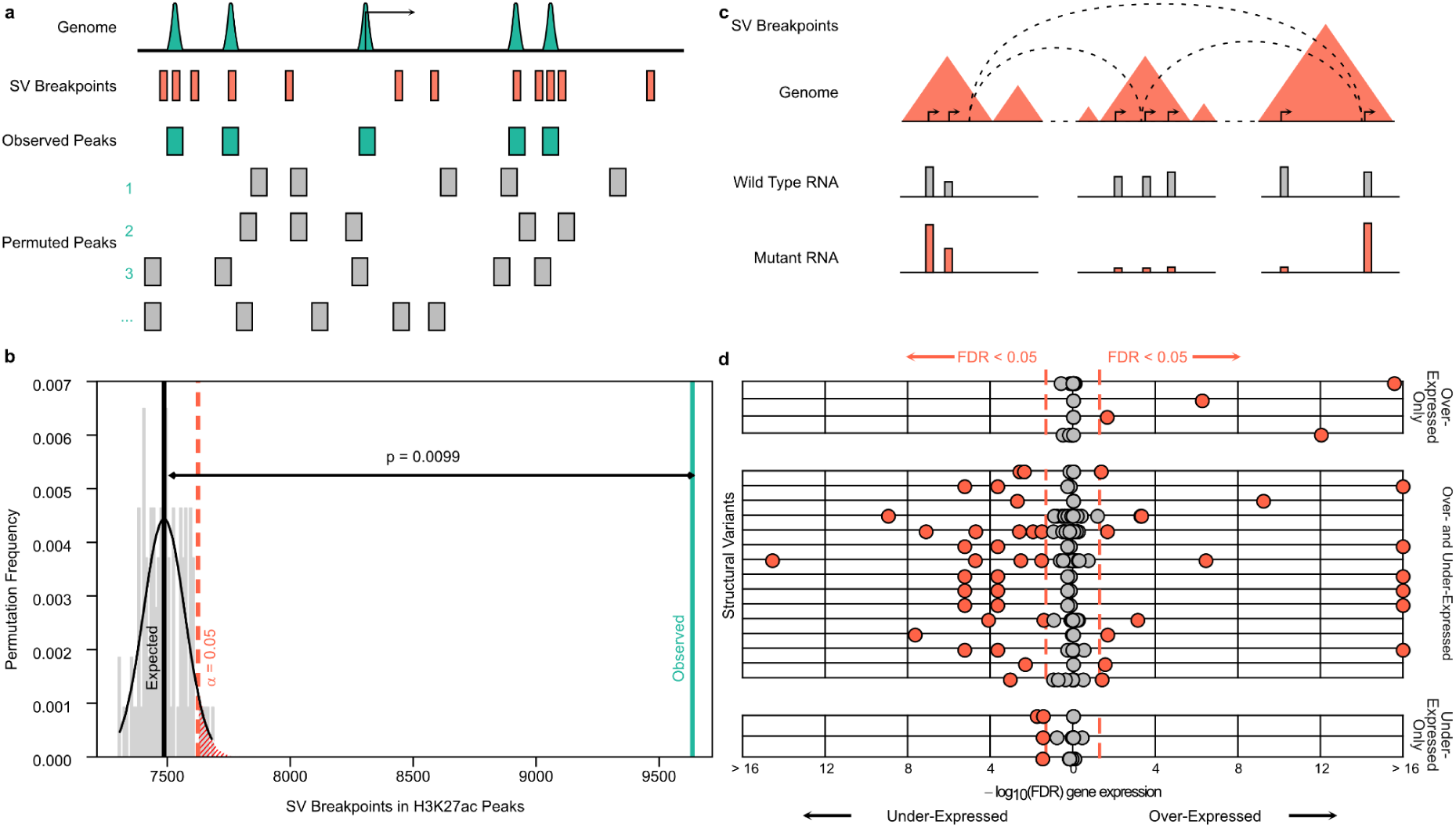
SV breakpoints are enriched in active CREs and repeatedly alter the expression of multiple genes. **a.** Schematic of permutation testing for the overlap between SV breakpoints in all CPC-GENE prostate tumours and the catalogue of active CREs in the 12 tumour samples in this study. **b.** Histogram of permutation test results in grey. The vertical black and green bars refer to the expected and observed overlap of SV breakpoints and CREs, respectively. P-value is obtained from the permutation test, *n* = 100. **c.** Schematic of how the expression of genes within TADs containing SV breakpoints are compared between mutant and wild type tumours are compared. **d.** Scatterplot of FDR values obtained from differential gene expression analysis as outlined in **c**. Red dots are differentially expressed genes (FDR < 0.05), grey dots are genes not differentially expressed between the mutant and wild type tumours.

The fusion of *PMEPA1* and *ZNF516* is an example of CRE hijacking resulting in opposite differential gene expression. Specifically, the fusion results in the *PMEPA1* promoter being hijacked to the 5’ end of the *ZNF516* gene. This is concomitant with the over-expression of *ZNF516* and under-expression of *PMEPA1* (Figure 6a-c). In addition to hijacking the *PMEPA1* promoter to the *ZNF516* gene, this fusion also coincides with gains in H3K27ac over the *ZNF516* gene body and of H3K27ac histone hypoacetylation over the 3’ end of *PMEPA1*’s gene body. This mirrors the creation of a Cluster Of Regulatory Elements (COREs) reported for the T2E fusion, reflective of new CREs enabling *ERG* over-expression and the concomitant under-expression of *TMPRSS2* (Supplementary Figure 7) ^42, 51, 52^. CRE hijacking is also observed with inter-chromosomal rearrangements such as seen at the SV connecting chromosomes 7 and 19, creating 2 fusion products (termed C2B and B2C; Figure 6d). This SV separates the 3’ end of *BRAF* from its promoter and upstream enhancers on chromosome 7 (C2B; Figure 6d), fusing it to the 3’ end of *CYP4F11* (Figure 6e). Focal chromatin interactions between *BRAF* and multiple active CREs are only observed in the fusion on chromosome 19 (Figure 6e). Using matched RNA-seq data, we observe an estimated 5 fold increase in expression for the 3’ exons of *BRAF* in the mutated tumour compared to others (fold-change = 4.976, FDR = 0.0181; Figure 6f). Collectively, over-expression of the oncogenes, such as *ERG* and *BRAF,* and suppression of the tumour suppressor *PMEPA1* demonstrates the disease-relevant effects of CRE hijacking mediated by SVs in primary prostate cancer resulting in changes to focal chromatin interactions, and that these effects overshadow the effect on higher-order topology in primary prostate cancer.

**Figure 6.**
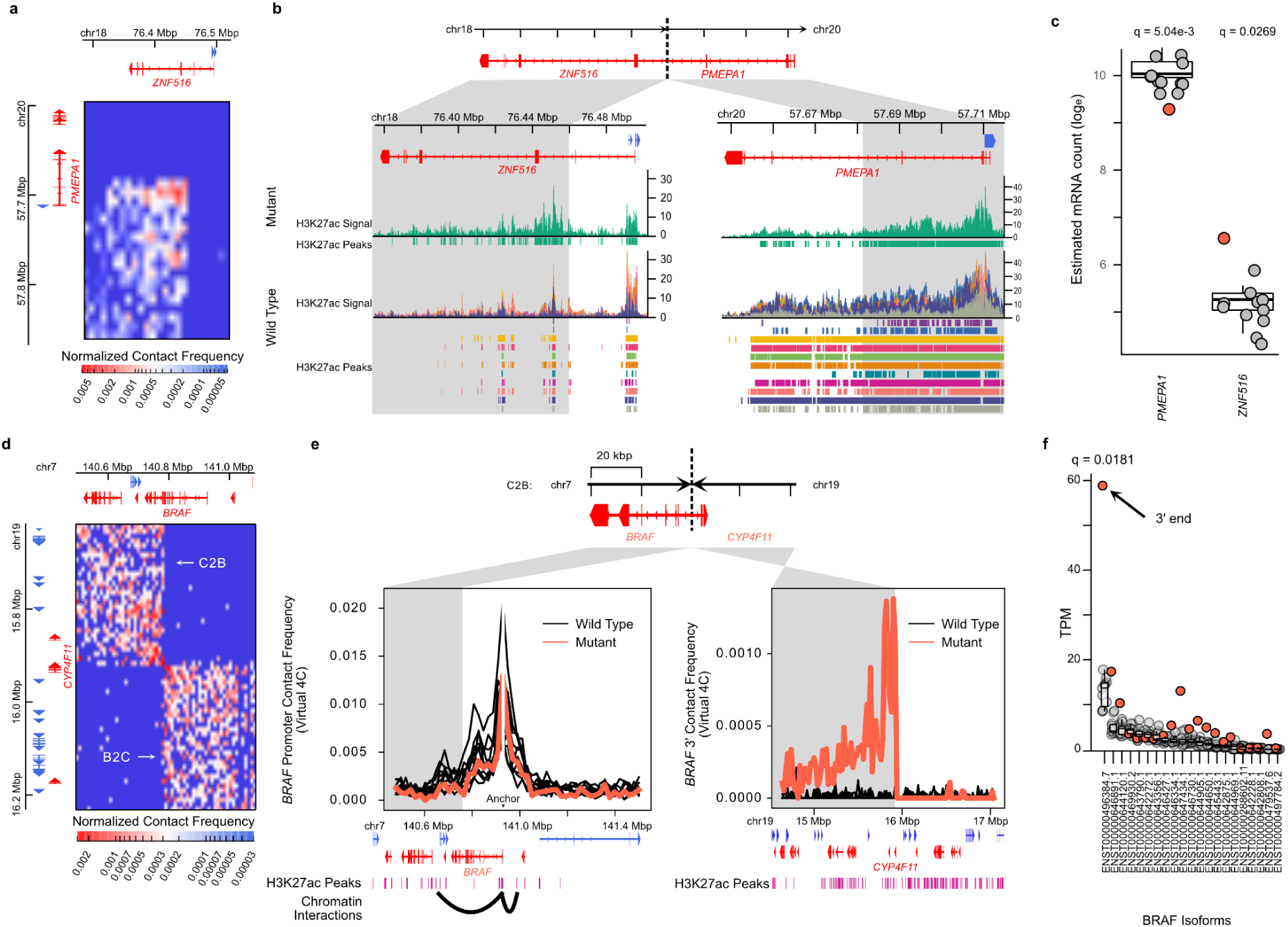
SVs altering gene expression by rewiring focal chromatin interactions. **a**. The contact matrix of the deletion between *PMEPA1* and *ZNF516.* **b**. Genome tracks of H3K27ac ChIP-seq signal around the *ZNF516* and *PMEPA1* genes with the rearrangement. Grey regions are loci brought into contact by the SV. **c**. Gene expression of *PMEPA1* and *ZNF516* in all tumour samples. Boxplots represent the first, second, and third quartiles of wild type patients (grey dots). Red dots are the gene expression for the mutated patient. **d**. The contact matrix of an inter-chromosomal break between chromosome 7 and chromosome 19. **e**. Contact frequencies of the *BRAF* promoter on chromosome 7 (left) and the 3’ end of *BRAF* on chromosome 19 (right). SV-associated contacts between the 3’ end of *BRAF* on chromosome 19 (right) are focally enriched at H3K27ac peaks downstream of *CYF4P11*. Bar plot of SVs categorized by how differentially expressed genes altered.

## Discussion

Genetic alterations that subvert the higher-order chromatin organization to allow for aberrant focal interactions may be more common in cancer than previously recognized. In this work we demonstrated that CRE hijacking by SVs is often associated with opposing gene expression changes at SV antipodes, whereby genes on one flank of the breakpoint are upregulated while genes on the other flank are repressed. Complex SVs, such as chromoplexy and chromothripsis, are found in numerous cancer types ^14, 38^, providing many opportunities for widespread effects on gene expression and CRE hijacking. This is in addition to many known cancer drivers that alter CRE interactions, including the *AR* and *FOXA1* enhancer amplifications in primary and metastatic prostate tumours ^19, 20, 34, 36, 42, 53^. More recent findings also fit this model, such as accumulation of extrachromosomal circular DNA activating oncogenes that would otherwise be constrained by chromatin topology ^54–57^. These insights stress the importance of investigating all ends of an SV to assess the biological impact of these mutations on the *cis*-regulatory landscape as a whole, as opposed to focusing on CREs or SV breakpoints as single entities.

Changes to the three-dimensional genome reported in disease onset or development are often inferred from alterations in TAD boundaries ^9, 28^. For instance, CTCF activity is targeted by somatic mutations that enrich at its binding sites in colorectal, oesophageal, and liver cancers ^26, 58^. Furthermore, gains in DNA methylation at CTCF binding sites are linked to altered TAD structures in gliomas ^7^. In primary PCa 97% of differentially methylated regions genome-wide in primary PCa are losses of DNA methylation ^59, 60^, an epigenetic process previously shown to have limited impact on CTCF chromatin binding ^61^. This suggests that aberrant CTCF binding at TAD boundaries is not a hallmark of prostate oncogenesis. Our observation of stable chromatin organization supports this model. Notably, stable TAD structures observed in these primary tissues contrast previous reports of chromatin organization in cell lines derived from prostate cells ^44, 62^, highlighting the necessity of low-input protocols and primary tissues ^40^. Our findings further support recent reports of shared higher-order chromatin organization among phenotypically distinct cell types in model organisms ^2, 28, 32, 63–65^. Taken together, this body of evidence suggests that large disruptions to TADs and compartments may constrain the transformation of normal to cancer cells or the divergent subtyping within prostate tumours. Instead, changes to focal chromatin interactions seem to reflect alterations in the genetic architecture leading to cancer development. Investigating these focal chromatin interactions may provide insights on the relationship between CREs, such as between enhancers and their target gene promoter ^66, 67^, to better understand the etiology of disease.

In conclusion, by bypassing technical limitations to characterize the three-dimensional genome organization across benign and tumour prostate tissue ^40^, our work reveals the predominant stable nature of genome topology across prostate oncogenesis. Instead, alterations to discrete chromatin interactions populate the PCa genome. These impact the function of CREs, such as we report for SV-mediated CRE hijacking events. Considering the contribution of SVs across human cancers ^68^, our collective work presents a framework inclusive of genetics, chromatin state, and three-dimensional genome organization to understand the genetic architecture across individual primary tumours.

## Materials & Methods

### Patient Selection Criteria

Patients were selected from the CPC-GENE cohort of Canadian men with indolent PCa, Gleason scores of 3+3, 3+4, and 4+3. All primary human material was obtained with informed consent with approval of our institutional research ethics board (UHN 11-0024). The intersection of previously published data for whole genome sequencing ^33^, RNA abundance ^41^, and H3K27ac ChIP-seq ^42^ led to 25 samples having data for all assays. 11 of these tested positive for ETS gene family fusions (T2E status), and 14 without. To accurately represent the presence of this subtype of PCa in the disease generally, and to ensure minimum read depths required to perform accurate analysis on chromatin conformation data, we selected approximately half of these remaining samples (6 T2E+ and 6 T2E-).

### Patient Tumour in situ low-input Hi-C Sequencing

We followed the general *in situ* low input Hi-C (Low-C) protocol from Díaz *et al.* ^40^, with our own re-optimization for solid tumour tissue sections. It is worth noting that throughout the protocol, the pellet would be hardly visible and would require careful pipetting. The specific modifications of the protocol are described below.

#### Tumour Tissue Preparation

Twelve cryopreserved-frozen PCa tumour tissue specimens were obtained from primary PCa patients as part of the Canadian PCa Genome Network (CPC-GENE) effort ^33^. Informed consent was obtained from all patients with REB approval (UHN 11-0024). These tumour specimens were sectioned into 10 µm sections. Sections before and after the sections used for Hi-C were stained with hematoxylin and eosin (H&E) and assessed pathologically for ≥ 70% PCa cellularity. The percentage of infiltrating lymphocytes was also estimated by pathological assessment to be ≤3%. Stratification into *TMPRSS2-ERG* (T2E)-positive or T2E-negative was determined through either whole-genome sequencing detection of the rearrangement, immunohistochemistry, or mRNA expression microarray data ^33^.

#### Normal Tissue Preparation

Five snap-frozen prostate tumour-adjacent normal tissue specimens were obtained. Informed consent was obtained from all patients with REB approval (UHN 11-0024). Tissue specimens were sectioned into 5, 10, and 20 µm sections. Sections used for Hi-C and RNA-seq were stained with H&E and assessed pathologically for ≥ 60% prostate glandular cellularity.

#### Fixation and Lysis

One or two sections (consecutive; depending on surface area) for each patient were thawed and fixed by adding 300 µL of 1% formaldehyde in PBS directly onto the tissue sample, followed by a 10-minute incubation at room temperature (RT) (Supplementary Figure 1a). The formaldehyde was quenched by adding 20 µL of 2.5M glycine to the sample reaching a final concentration of 0.2M followed by 5 minutes of incubation at RT. The samples were then washed three times with 500 µL cold PBS and scraped off the microscope slide with a scalpel into 1.5 mL centrifuge tube containing 250 µL of ice-cold Low-C lysis buffer (10 mM Tris-Cl pH 8.0, 10 mM NaCl, 0.2% IGEPAL CA-630 (Sigma-Aldrich)) supplemented with protease inhibitor. The samples were then mixed thoroughly by gentle pipetting and left on ice for 20 minutes with intermittent mixing. Upon lysis, the samples were snap-frozen with liquid nitrogen and stored at -80 °C until processing the next day. As a note, stagger fixation times when processing multiple samples to prevent needless rush and chance of under/over-fixation.

#### Enzyme Digestion and Overhang Fill-In

The samples stored at -80 °C were thawed on ice and spun down at 300 x g for 5 minutes at 4 °C. The samples were then resuspended in 125 µL of ice-cold 10X NEB2 Buffer (New England Biolabs), and again spun down at 13,000 x g for 5 minutes at 4 °C. The pellet was then resuspended in 25 µL of 0.4% SDS and incubated at 65 °C for 10 minutes without agitation for permeabilization. To quench the SDS, 10% Triton X-100 in water (12.5 µL + 75 µL water) was then added to the samples and incubated at 37 °C for 45 minutes at 650 rpm. For enzymatic digestion, 35 µL of 10X NEB2.1 buffer (New England Biolabs) was added to each sample, follow by the addition of 50 U of MboI and 90 minutes incubation at 37 °C with gentle agitation (add 30 U first, incubate 45 minutes, followed by the addition of another 20 U and another 45 minutes of incubation). Upon digestion, the MboI enzyme was inactivated by incubating at 62 °C for 20 minutes. The overhangs generated by the MboI enzyme was then filled-in by adding a mix of dNTPs and DNA Polymerase I Klenow Fragment directly to each sample (10 µL of 0.4 mM biotin-14-dCTP, 0.5 µL of 10 mM dATP, 0.5 µL of 10 mM dGTP, 0.5 µL of 10 mM dTTP, 4 µL of 5U/µL DNA Polymerase I Klenow Fragment). The samples were then mixed by gentle pipetting followed by incubation at 37 °C for 90 minutes with gentle agitation.

#### Proximity Ligation and Decrosslinking

Upon overhang fill-in, each sample was subject to proximity ligation through the addition of 328.5 µL water, 60 µL of 10X T4 DNA Ligase Buffer (ThermoFisher Scientific), 50 µL of 10% Triton X-100, 6 µL of 20 mg/mL BSA (New England Biolabs) and 3.5 µL of 5 Weiss U/µL T4 DNA Ligase (ThermoFisher). The samples were mixed through gentle pipetting and incubated at RT (20-22 °C) with rotation for 4 hours. The samples were then spun down at 13,000 x g for 5 minutes at RT and resuspended in 250 µL of Extraction Buffer (50 mM Tris-Cl pH 8.0, 50 mM NaCl, 1 mM EDTA, 1% SDS) upon removal of supernatant. Next, 10 µL of 20 mg/mL Proteinase K (New England Biolabs) was added to each sample and incubated at 55 °C for 30 minutes at 1,000 rpm. Then 65 µL of 5 M NaCl was added to each sample and incubated at 65 °C at 1,000 rpm overnight.

### DNA Extraction

Phenol-chloroform extraction columns were spun down at 17,000 x g for 1 minute at 4 °C to get gel down to the bottom of the tube. The samples incubated overnight were then added to the column. Next, an equal volume (∼325 µL) of phenol-chloroform-isoamyl alcohol mixture (25:24:1) (Sigma) was also added to the column. The column was then inverted for thorough mixing and spun down at 17,000 x g for 5 minutes at 4 °C. The surface layer on top of the gel upon spinning contains the sample and is transferred to a clean 1.5 mL tube (∼325 µL). Each sample was mixed with 31.5 µL of 3M sodium acetate, 2 µL of GlycoBlue (ThermoFisher Scientific), and 504 µL of 100% ethanol for DNA precipitation. The samples were inverted several times for mixing and incubated at -80 °C for 20 minutes, followed by a centrifuge spin at 17,000 x g for 45 minutes at 4 °C. The supernatant was carefully discarded and the pellet was washed with 800 µL of ice-cold 70% ethanol followed by a centrifuge spin at 17,000 x g for 5 minutes at 4 °C. The supernatant was then discarded and the tube was air-dried until no traces of ethanol was left prior to dissolving the DNA pellet with 30 µL of Elution Buffer (Qiagen PCR Clean-Up Kit). 1 µL of RNase A (ThermoFisher Scientific) was added to each sample followed by incubation at 37 °C for 15 minutes. A mix of 5 µL of 10X NEB2.1 buffer (New England Biolabs), 1.25 µL of 1 mM dATP, 1.25 µL of 1 mM dCTP, 1.25 µL of 1 mM dGTP, 1 mM of dTTP, 0.5 µL of 10 mg/mL BSA, 5 µL of water, 3.5 µL of 3 U/µL T4 DNA Polymerase (New England Biolabs) was added to each sample. The samples were mixed thoroughly by gentle pipetting, and then incubated at 20 °C for 4 hours.

#### Fragmentation and Biotin Pull-down

70 µL of water was added to each sample bringing total volume up to 120 µL, and the samples were transferred into Covaris sonication tubes. The samples were then sonicated using Covaris M220 sonicator to attain 300-700 bp fragments. For biotin pull-down using a magnetic rack, 30 µL of Dynabeads MyOne Streptavidin C1 beads (Life Technologies) for each sample was washed once with 400 µL of 1X B&W buffer + 0.1% Triton X-100. The beads were then resuspended in 120 µL of 2X B&W buffer and transferred to the 120 µL of sample (1:1 ratio). The sample was then incubated with gentle rotation at RT for 20 minutes. The supernatant was discarded and the beads were resuspended with 400 µL of 1X B&W buffer + 0.1% Triton X-100 followed by a 2-minute incubation at 55 °C with mixing. The wash was repeated once more, then resuspended in 400 µL of 1X NEB2 buffer (New England Biolab).

#### Library Preparation and Size Selection

The beads containing the Hi-C samples were separated on a magnetic rack to remove the supernatant. The beads were then resuspended in a total volume of 10 µL for library preparation using the SMARTer ThruPLEX DNA-seq library preparation kit (Takara Biosciences) per manufacturer’s protocol with an adjustment on the last step, a PCR reaction for library amplification. Upon reaching that step, the reaction was carried out on a regular PCR for two cycles to amplify the Hi-C samples off the streptavidin beads. Next, the samples were transferred onto a new tube where 20X SYBR was added. The samples were then subject to real-time qPCR and pulled out from the qPCR machine mid-exponential phase. Ultimately, this is done to reduce PCR duplication rates, a huge limitation for low-input Hi-C protocols. The Hi-C libraries were then double size-selected for 300-700 bp using Ampure XP beads and sent for BioAnalyzer analysis prior to sequencing.

### Hi-C Sequencing and Data Pre-processing

### Sequencing

The Hi-C libraries for each tumour sample were sent for shallow paired-end 150 bp sequencing (∼10-15 million reads per sample) on a NextSeq 500. Upon confirming library quality and low duplication rates (< 2%), samples were sent for deep paired-end 150 bp sequencing with the aim of 800 million raw read pairs per sample on NovaSeq 6000.

#### Sequence alignment and Hi-C artefact removal

Paired-end FASTQ files were pre-processed with HiCUP (v0.7.2) ^69^. Reads were truncated at MboI ligation junction sites prior to alignment with ‘hicup_digester’. Each mate was independently aligned to the hg38 genome and were then paired and assigned to MboI restriction sites by ‘hicup_map’. ‘hicup_map’ uses Bowtie2 (v2.3.4) ^70^ as the underlying aligner which has the following parameters: ‘--very-sensitive --no-unal --reorder’. Reads that reflect technical artefacts were filtered out with ‘hicup_filter’. Duplicate reads were removed with ‘hicup_deduplicator’.

Reads that came from different sequencing batches were then aggregated for each tumour sample at this stage using ‘sambamba mergè (v0.6.9) ^71^. This resulted in an average of 1.12 x 10^9^ read per tumour sample (Supplementary Table 2).

#### Contact matrix generation and balancing

Aggregated binary alignment map (BAM) files were converted to the pairs format using pairtools (v0.2.2) ^72^ and then the cooler format using the cooler package (v0.8.5) ^73^. The pairs files were generated with the following command: ‘pairtools parse -c {genome} --assembly hg38 -o {output_pairs} {input_bam}’. The cooler files were generated at an initial matrix resolution of 1000 bp with the following command: ‘cooler cload pairs --assembly hg38 -c1 2 -p1 3 -c2 4 -p2 5 {genome}:1000 {input_pairs} {output_cooler}’.

The raw contact matrices stored in the cooler file format were balanced using cooler’s implementation of the ICE algorithm ^74^ using the ‘cooler balancè command. Contact matrices at different resolutions were created with the ‘cooler zoomify’ command.

### Hi-C Data Analysis

#### TAD identification

Contact matrices were binned at a resolution of 40 kbp. To remove sequencing depth as a confounding factor, contact matrices for all samples were first downsampled to match the sequencing depth of the shallowest sample. For comparisons including cell lines, this was 120x10^6^ contacts. For comparisons only involving primary samples, this was 300x10^6^ contacts. This was achieved with Cooltools (v0.3.2) with the following command: ‘cooltools random-sample -c 120000000 {input}::/resolutions/40000 {output}’.

TADs were identified using TopDom ^75^ on the downsampled, ICE-normalized contact matrices. To identify domains at multiple length scales, similar in concept to Artamus’ gamma parameter ^76^, TopDom was run multiple times per sample, with the window size parameter set at values between 3 and 40, inclusive (corresponding to 120 kbp and 1.6 Mbp). The lower bound for the window size parameter allowed for the identification of domains multiple megabases in size at the upper end and domains < 100 kbp at the lower end without being dominated by false calls due to sparsity of the data. Despite TopDom being more resistant to confounding by sequencing depth than other TAD calling tools ^77^, biases in boundary persistence were evident between samples of different sequencing depth. Downsampling contact matrices to similar depths resolved these biases.

Given the stochasticity of Hi-C sequencing, boundaries called at one window size may not correspond to the exact same location at a different window size. To attempt to resolve these different boundary calls and leverage power from multiple window sizes, boundaries for a given patient were considered at all window sizes. Boundaries within one bin (40 kbp) of each other and called at different window sizes were marked as conflicting calls. If only two boundaries were in conflict and all the window sizes where the first boundary was called are smaller than the window sizes where the second boundary was called, the second boundary was selected since larger smoothing windows are less sensitive to small differences in contact counts. If only two boundaries were in conflict but there is no proper ordering of the window sizes, the boundary that was identified most often between the two was selected. If three boundaries are in conflict, the middle boundary was selected. If four or more boundaries were in conflict, the boundary that was identified most often was selected.

To determine the maximum window size for TAD calls, TAD calls were compared across window sizes for the same patient using the BPscore metric ^78^. TAD calls are identical when the BPscore is 0, and divergent when 1. The cut-off window size for a single patient was determined when the difference between TAD calls at consecutive window sizes was < 0.005, twice in a row. The maximum window size was determined by the maximum window size cut-off across all samples in a comparison. For comparisons involving only primary samples, the maximum window size was determined to be w = 20 x 40 kbp. For comparisons involving cell lines, this was w = 32 x 40 kbp.

The persistence of a TAD boundary was calculated as the number of window sizes where this region was identified as a boundary.

#### Sample clustering by TADs

Using the TAD calls at the window size w = 32 x 40 kbp, the similarity between samples was calculated with BPscore. The resulting matrix, containing the similarity between any two samples, was used as the distance matrix for unsupervised hierarchical clustering with Ward.D2 linkage.

#### Compartment identification

Contact matrices were binned at a resolution of 40 kbp, similarly to TAD identification. To remove sequencing depth as a confounding factor, contact matrices for all samples were first downsampled to match the sequencing depth of the shallowest sample. Contact matrix eigenvectors were calculated with Cooltools. To standardize the sign of each eigenvector, the GC content of the reference genome, binned at 40 kbp, was used as a phasing reference track. This reference track was calculated with the ‘frac_gc’ function from the Bioframe Python package (v0.0.12) ^79^. The first eigenvector was used to identify compartments with the following command: ‘cooltools call-compartments --bigwig --reference-track gc-content-phase.bedGraph -o {output}{input}’.

#### Identification of significant chromatin interactions

Chromatin interactions were identified in all 17 primary samples with Mustache (v1.0.2) ^80^. Using the Cooler files from above, Mustache was run on the ICE-normalized 10 kbp contact matrix for each chromosome with the following command: ‘mustache -f {input} -r 10000 -ch {chromosome} -p 8 -o {output}’. Interaction calls on each chromosome were merged for each sample to create a single table of interaction calls across the entire genome.

To account for variances in detection across samples and to identify similarly called interactions across samples, interaction anchors were aggregated across all samples to form a consensus set. Interaction anchors were merged if they overlapped by at least 1 bp. Interaction anchors for each sample were then mapped to the consensus set of anchors, and these new anchors were used in all subsequent analyses.

#### Chromatin interaction saturation analysis

To estimate the detection of all chromatin interactions across all samples, a nonlinear regression on an asymptotic model was performed. This is similar in method to peak saturation analysis used to assess peaks detected in ChIP-seq experiments from a collection of samples ^42^. Bootstrapping the number of unique interactions detected in a random selection of *n* samples was calculated for *n* ranging from 1 to 17. 100 iterations of the bootstrapping process were performed. An exponential model was fit against the mean number of unique interactions detected in *n* samples using the ‘nls’ and ‘SSaymp’ functions from the stats R package (v3.6.3). The model was fit to the following equation:

Where is the mean number of chromatin interactions for a given number of samples, is the asymptotic limit of the total number of mean detected interactions, is the response for, and is the rate constant. The estimated fit was used to predict the number of samples required to reach 50%, 90%, 95%, and 99% saturation of the asymptote (Supplementary Figure 3c).

#### Structural variant breakpoint pair detection

Breakpoint pairs for each patient were called on the merged BAM files using ‘hic_breakfinder’ (commit 30a0dcc6d01859797d7c263df7335fd2f52df7b8) ^27^. Pre-calculated expected observation files for the hg38 genome were downloaded from the ‘hic_breakfinder’ GitHub repository on Jul 24, 2019, as per the instructions. Breakpoints were explicitly called with the following command: ‘hic_breakfinder --bam-file {BAM} --exp-file-inter inter_expect_1Mb.hg38.txt --exp-file-intra intra_expect_100kb.hg38.txt --name {Sample ID} --min-1kb’.

For the T2E fusion, only one patient had the deletion identified by hic_breakfinder with default parameters (CPCG0336). Difficulties identifying SVs with hic_breakfinder have been previously noted ^47^. After adjusting the detection threshold, we were able to identify the fusion in other samples. To ensure the T2E+ tumours were effectively stratified for future analyses, the fusion was annotated using the same coordinates for the other T2E+ samples. No other additions to breakpoint calls were made. Certain breakpoints that appeared to be artefacts were removed, as described below.

#### Structural variant annotation and graph construction

The contact matrix spanning 5 Mbp upstream and downstream around the breakpoint pairs were plotted and annotated according to previously published heuristics (Supplementary Figure 4 for ^27^). Breakpoint pairs that were nearby other breakpoints or did not match the heuristics in this figure were labelled as “unknown”. These annotations were matched against the annotations identified from the previously published whole genome sequencing structural variants ^33^.

Breakpoint pairs matching the following criteria were considered as detection artefacts and were ignored.

1. At least one breakpoint was > 1 Mbp
2. At least one breakpoint was surrounded by empty regions of the contact matrix
3. At least one breakpoint corresponded to a TAD or compartment boundary shared across all samples that lacked a distinct sharp edge that is indicative of a chromosomal rearrangement To identify unique breakpoints that were identified in multiple breakpoint pairs, breakpoints that were within 50 kbp of each other were considered as possibly redundant calls. This distance was considered as the resolution of the non-artefactual calls is 100 kbp. Plotting the contact matrix 5 Mbp around the breakpoint, breakpoints calls were considered the same breakpoint if the sharp edge of each breakpoint was equal to within 5 kbp.

Similar in concept to the ChainFinder algorithm ^38^, we consider each breakpoint as a node in a graph. Nodes are connected if they are detected as a pair of breakpoints by ‘hic_breakfinder’. Simple structural variants are connected components in the breakpoint graph containing only two nodes, and complex variants those with greater than two nodes. A visual representation of these graphs can be found in Supplementary Figure 4b.

#### Determination of structural variant breakpoints altering TAD boundaries

Patients are assigned into one of two groups using hierarchical clustering (complete linkage) with the matrix of pairwise BPscore ^78^ values as a distance matrix. If the clustering equals the mutated samples from the non-mutated samples (i.e., the clustering matches the mutation status in this locus), then the local topology was classified as “altered” because of the SV.

#### Virtual 4C

Two parts of the *BRAF* gene were used as anchors for virtual 4C data: the promoter region (1500 bp upstream, 500 bp downstream of the TSS) and the entire gene downstream of the breakpoint. Contact frequencies from the ICE-normalized, 20 kbp contact matrices were extracted, with the rows as the bins containing the anchor and the columns as the target regions (the x-axes in Figure 6e). The row means were calculated to produce a single vector where each element is the average normalized contact frequency between the anchor of interest and the distal 20 kbp bin. These vectors were plotted as lines in Figure 6e.

### Patient Tumour Tissue H3K27ac ChIP-seq

ChIP-seq against H3K27ac was previously published for these matching samples in ^42^. Sequencing data was processed similarly to the previous publication of this data ^42^; however, the hg38 reference genome was used instead of hg19.

#### Sequence alignment

FASTQ files from single-end sequencing were aligned to the hg38 genome using Bowtie2 (v2.3.4) with the following parameters: ‘-x {genome} -U {input} 2> {output_report} | samtools view -u > {output_bam}’. For FASTQ files from paired-end sequencing, only the first mate was considered and reads were aligned with the following parameters: ‘-x {genome} -U {input} -3 50 2> {output_report} | samtools view -u > {output_bam}’. This ensured that all H3K27ac ChIP-seq data had the same format (single-end) and length (52 bp) before alignment to mitigate possible differences in downstream analyses due to different sequencing methods. Duplicate reads were removed with sambamba (v0.6.9) via ‘sambamba markdup -r’ and were then sorted by position using ‘sambamba sort’.

#### Peak calling

Peak calling was performed using MACS2 (v2.1.2) ^81^ with the following command: ‘macs2 callpeak -g hs -f BAM -q 0.005 -B -n {output_prefix} -t {seq_chip} -c {seq_input}’. ENCODE hg38 blacklist regions were then removed from the narrow peaks ^82^. Peaks calls are in Supplementary Table 8.

#### Differential acetylation analysis

Unique peak calls and deduplicated pull-down and control BAM files from tumour samples were loaded into R with the DiffBind package (v2.14.0) ^83^ using the DESeq2 (v1.26.0) ^84^ as the differential analysis model. 3 of the 12 samples had low quality peak calls compared to the other 9 and were not considered when calculating differential acetylation (CPCG0268, CPCG0255, and CPCG0336). We considered each unique breakpoint one at a time in the remaining 9 samples. Samples were grouped by their mutation status (i.e., a design matrix where the mutation status is the only covariate) and DiffBind’s differential binding analysis method was performed to identify all differentially acetylated regions between the two groups. Acetylation peaks outside of the TAD(s) overlapping the breakpoint were filtered out. Multiple test correction with the Benjamini-Hochberg FDR method was performed on all peaks after all breakpoints were considered, due to similar group stratifications depending on the breakpoint under consideration.

#### Structural variant breakpoint enrichment

Structural variant breakpoint coordinates from WGS data from the CPC-GENE cohort were obtained from the International Cancer Genome Consortium (structural somatic mutations from the PRAD-CA dataset, release 28). Breakpoint coordinates were lifted over to hg38 coordinates using the liftOver function from the rtracklayer R package (v1.46.0) ^85^. Permutation tests were performed with the regioneR R package (v1.18.0) ^86^, selecting randomized regions from the hg38 genome, excluding the ENCODE blacklist regions ^82^ and masked loci. 100 permutations were calculated and a one-sided permutation z-test was used to calculate statistical significance.

### Primary Tissue RNA Data Analysis

#### Tumour sample RNA sequencing

Total RNA was extracted for the CPC-GENE tumour samples as previously described ^41^. Briefly, total RNA was extracted with mirVana miRNA Isolation Kit (Life Technologies) according to the manufacturer’s instructions. RNA samples were sent to BGI Americas where it underwent QC and DNase treatment. For each sample, 200 ng of total RNA was used to construct a TruSeq strand-specific library with the Ribo Zero protocol (Illumina, Cat. #RS-122-2203). The libraries were sequenced on a HiSeq 2000 to a minimal target of 180 million, 2 x 100 bp paired-end reads.

#### RNA sequencing data pre-processing

RNA sequencing FASTQ files were pseudo-aligned to the hg38 genome using Kallisto (v0.46.1) ^87^ with the following command: ‘kallisto quant --bootstrap-samples 100 --pseudobam --threads 8 --index /path/to/GRCh38.idx --output-dir {output_dir} {input_R1.fastq.gz} {input_R2.fastq.gz}’.

#### Differential gene expression analysis

To assess whether SVs were associated with local gene expression changes, we considered each unique breakpoint one at a time. For each breakpoint, we compared the gene expression between the mutated and non-mutated tumour samples using Sleuth (v0.30.0) ^88, 89^ with a linear model where the mutation status was the only covariate. To reduce the chance of falsely identifying genes as differentially expressed, only genes located within the TADs (window size 20) containing breakpoints were considered. Fold-change estimates of each transcript were assessed for significance using a Wald test. Transcript-level p-values are combined to create gene-level p-values using the Lancaster aggregation method provided by the Sleuth package. Correcting for multiple tests was then performed with the Benjamini-Hochberg FDR correction for all genes that were potentially altered in the mutated sample(s).

### Data Availability

Whole genome and RNA sequencing data is available in the European Genome-Phenome Archive (EGA) under accession number EGAS00001000900. H3K27ac ChIP sequencing data is available in EGA under accession number EGAS00001002496. Hi-C sequencing data is under submission to the European Genome-Phenome Archive (EGAS00001005014). TADs and chromatin interactions are available in the Gene Expression Omnibus (GEO) archive under accession number GSE164347. Hi-C sequencing data from cell lines was obtained from GEO under expression number GSE118629 (22Rv1, RWPE1, and C4-2B cell lines) and from the 4D Nucleome under accession numbers 4DNFI6HDY7WZ (H1-hESC Rep 1), 4DNFITH978XV (H1-hESC Rep 2), 4DNFIT64Q7A3 (HAP-1 Rep 1), 4DNFINSKEZND (HAP-1 Rep 2), 4DNFIIV4M7TF (GM12878 Rep 1), and 4DNFIXVAKX9Q (GM12878 Rep 2). Processed data is available on CodeOcean (https://codeocean.com/capsule/5232537/tree).

### Code Availability

Code for data processing, analysis, and plotting can be found on CodeOcean (https://codeocean.com/capsule/5232537/tree).

## Author Contributions

S.Z., J.R.H., and M.L. conceptualized the study. S.Z. designed and conducted all the experiments with help from C.A. G.G., AND K.K. J.R.H. implemented all the computational and statistical approaches and analyses. R.H.-W. pre-processed the RNA-seq data from the primary tumours. Figures were designed by S.Z. and J.R.H. The manuscript was written by S.Z., J.H., and M.L with assistance from all authors. T.v.d.K., M.F., P.C.B., R.G.B., and M.L. supervised the study. M.L. oversaw the study.

## Supporting information

Supplementary Table 1 - Clinical Features

Supplementary Table 2 - Hi-C Sequencing Metrics

Supplementary Table 3 - TAD Statistics

Supplementary Table 4 - Individual TAD Calls

Supplementary Table 5 - Chromatin Interactions

Supplementary Table 6 - SV Breakpoint Pairs

Supplementary Table 7 - Simple and Complex SV Statistics

Supplementary Table 8 - H3K27ac ChIP-seq Peaks

## Acknowledgements

We thank all the Lupien lab members for their feedback, as well as Jesse Dixon for his support with hic_breakfinder and interpretation of structural variant calls. This work is supported by Prostate Cancer Foundation Canada, Ontario Institute for Cancer Research funded by the Government of Ontario, the Princess Margaret Cancer Foundation (R.G.B. and M.L.), Princess Margaret Cancer Centre Department of Surgical Oncology (M.F.), Princess Margaret Cancer Centre Genetics and Epigenetic Program (M.F. and M.L.), University of Toronto Department of Surgery Division of Urology (M.F.), Movember Foundation (RS2014-04 to M.L. and RS2014-01 to P.C.B.), the Radiation Medicine Program Academic Enrichment Fund (R.G.B.), Terry Fox Research Institute New Investigator Award (P.C.B.), Canadian Institute of Health Research (CIHR; FRN-153234 to M.L.) and New Investigator Award (P.C.B. and M.L.), Canadian Cancer Society Research Scientist Award (R.G.B.), Cancer Society Impact Award (P.C.B), Investigator Award from the Ontario Institute for Cancer Research (M.L. and P.C.B), and Movember Rising Star Award from PCa Canada (M.L. and P.C.B).

## Supplementary Figures

**Supplementary Figure 1.**
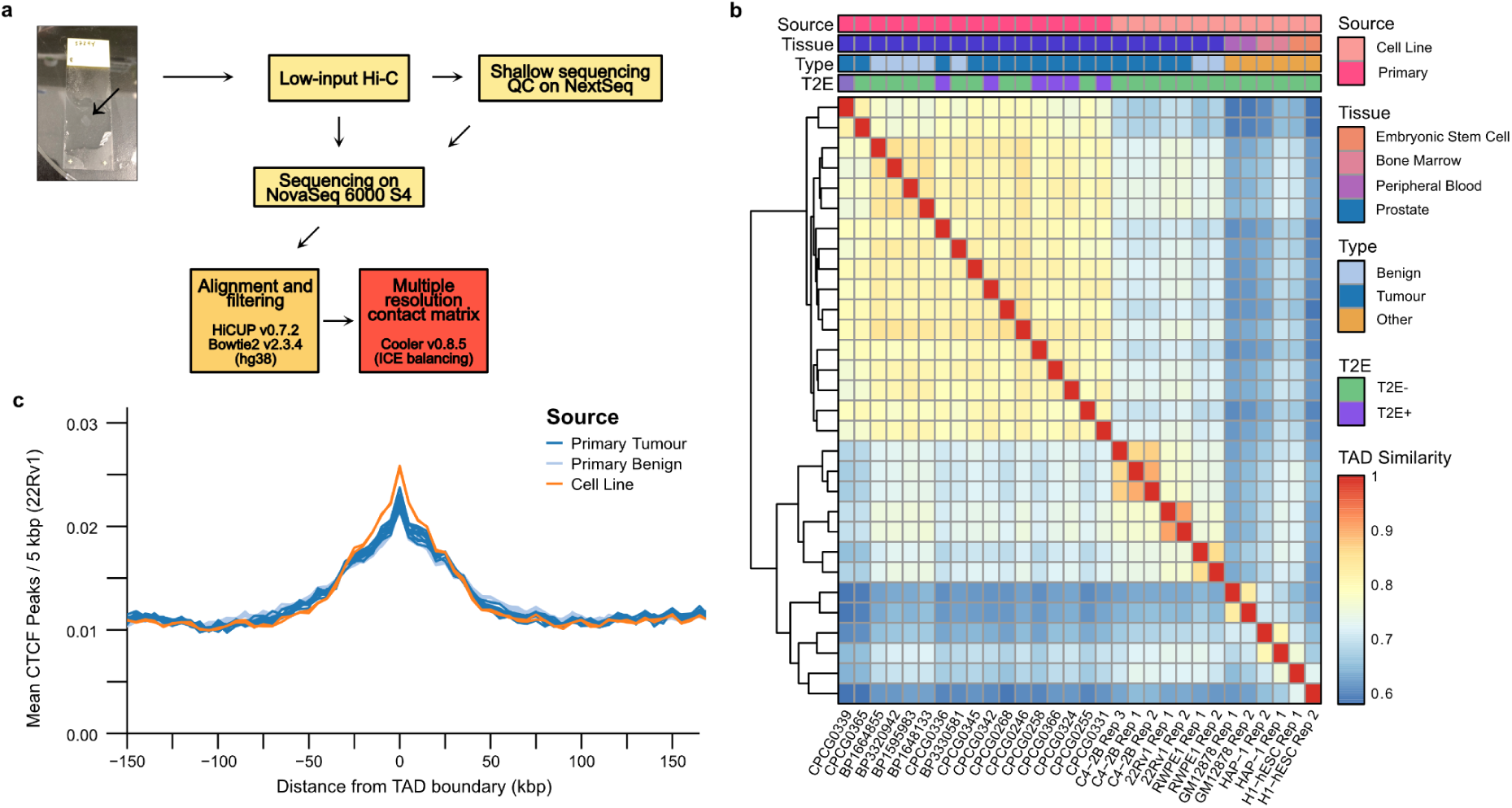
Sample processing and TAD similarity between samples. **a.** Schematic representation of the protocol and data pre-processing pipeline used in this study to obtain Hi-C sequencing data. **b.** Heatmap of TAD similarities between primary prostate samples, prostate cell lines, and non-prostate cell lines. Median similarity scores between TADs in primary prostate tissues and cell lines is 72.1%, 66.9% between prostate and non-prostate cell lines, and 63.5% between primary prostate and non-prostate lines. **c**. Local enrichment of CTCF binding sites from the 22Rv1 PCa cell line around TAD boundaries identified in the primary samples.

**Supplementary Figure 2.**
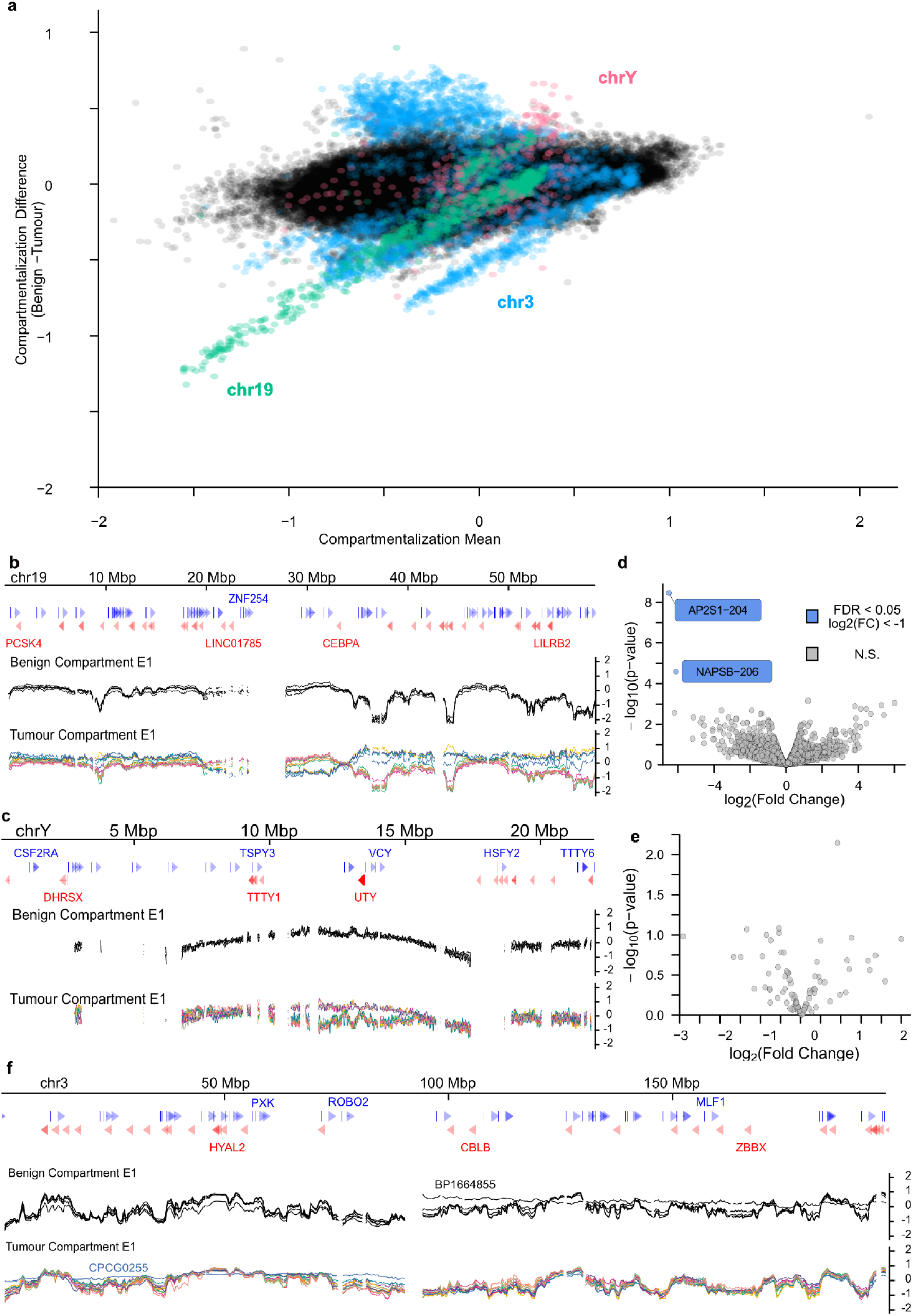
Compartmentalization changes in tumours is not associated with widespread differential gene expression. **a.** Bland-Altman plot of the mean compartmentalization score between tumour and benign samples. Chromosomes 3, 19, and Y are highlighted for their consistent deviation between the tissue types. **b-c.** Compartmentalization genome tracks across chromosomes 19 (**b**) and Y (**c**) in all primary samples. **d-e**. Volcano plot of differential transcript expression between the tumour samples with benign-like compartmentalization and altered compartmentalization in chromosomes 19 (**d**) and Y (**e**). Grey dots are transcripts without significant differential expression, blue dots are differentially expressed transcripts (FDR < 0.05) that are under-expressed in the altered compartment samples. **f.** Compartmentalization genome tracks across chromosome 3.

**Supplementary Figure 3.**
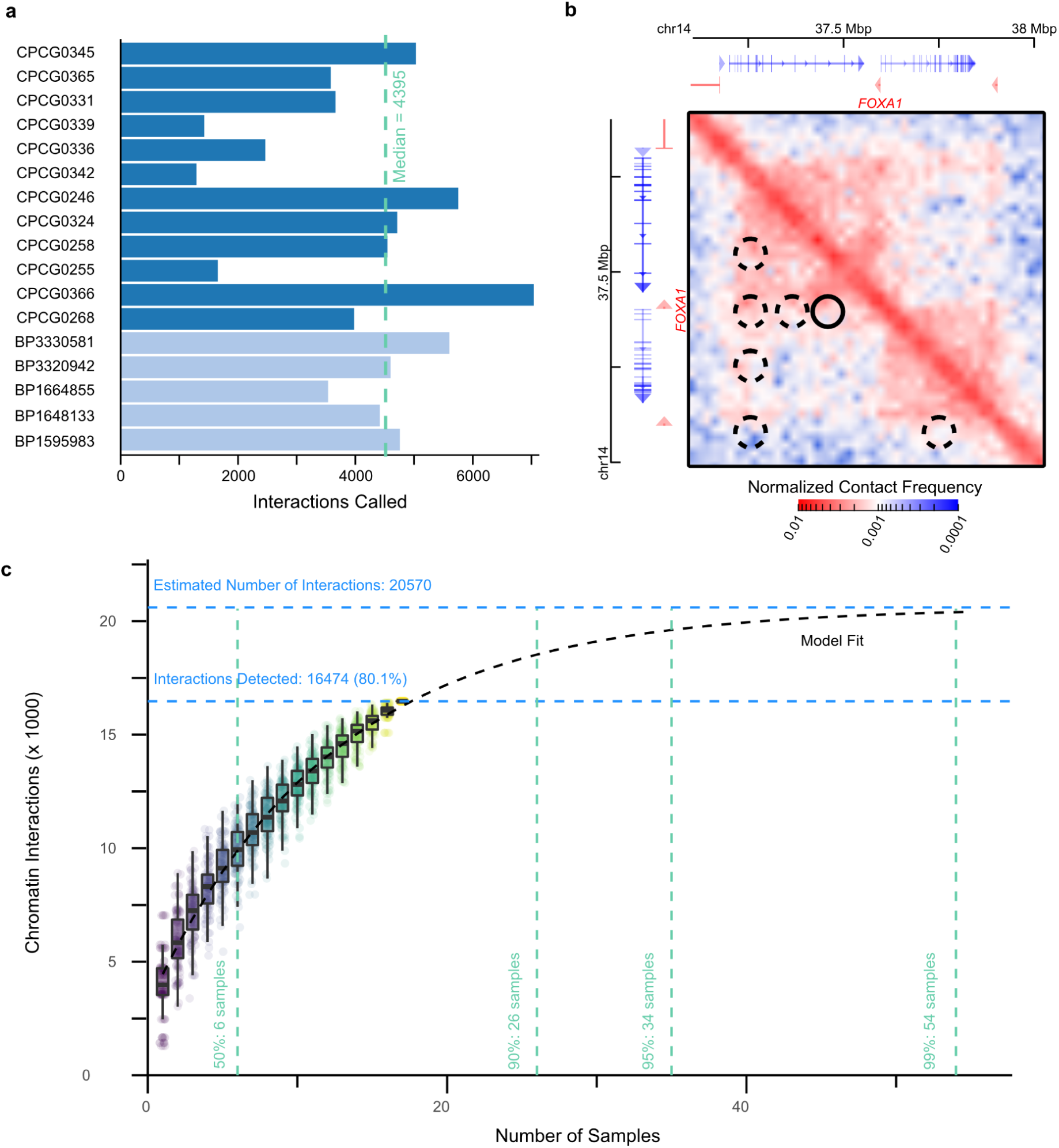
Characterization of chromatin interactions in benign and tumour tissue. **a.** Barplot of the number of significant chromatin interactions identified in each of the primary prostate samples. **b.** A snapshot of significant chromatin interactions called around the *FOXA1* gene. Identified interactions are highlighted as circles. The interaction marked by the solid border contains two CREs of *FOXA1* identified in REF^19^ (listed in that publication as CRE1 and CRE2). The interactions marked by the dashed border indicate regions of increased contact that may contain more distal CREs of *FOXA1*. **c.** Saturation analysis chromatin interactions detected in our cohort of prostate samples versus the theoretical estimation obtained through asymptotic estimation from bootstraps. Boxplots show the first, second, and third quartiles of the identified interactions across the bootstrap iterations. The dashed black line corresponds to the asymptotic model of estimated mean unique interactions obtained from an increasing number of samples. Horizontal blue dashed lines indicate the number of observed unique interactions and theoretical maximum. Vertical green dashed lines indicate the number of samples required to reach as estimated 50%, 90%, 95%, and 99% of the theoretical maximum.

**Supplementary Figure 4.**
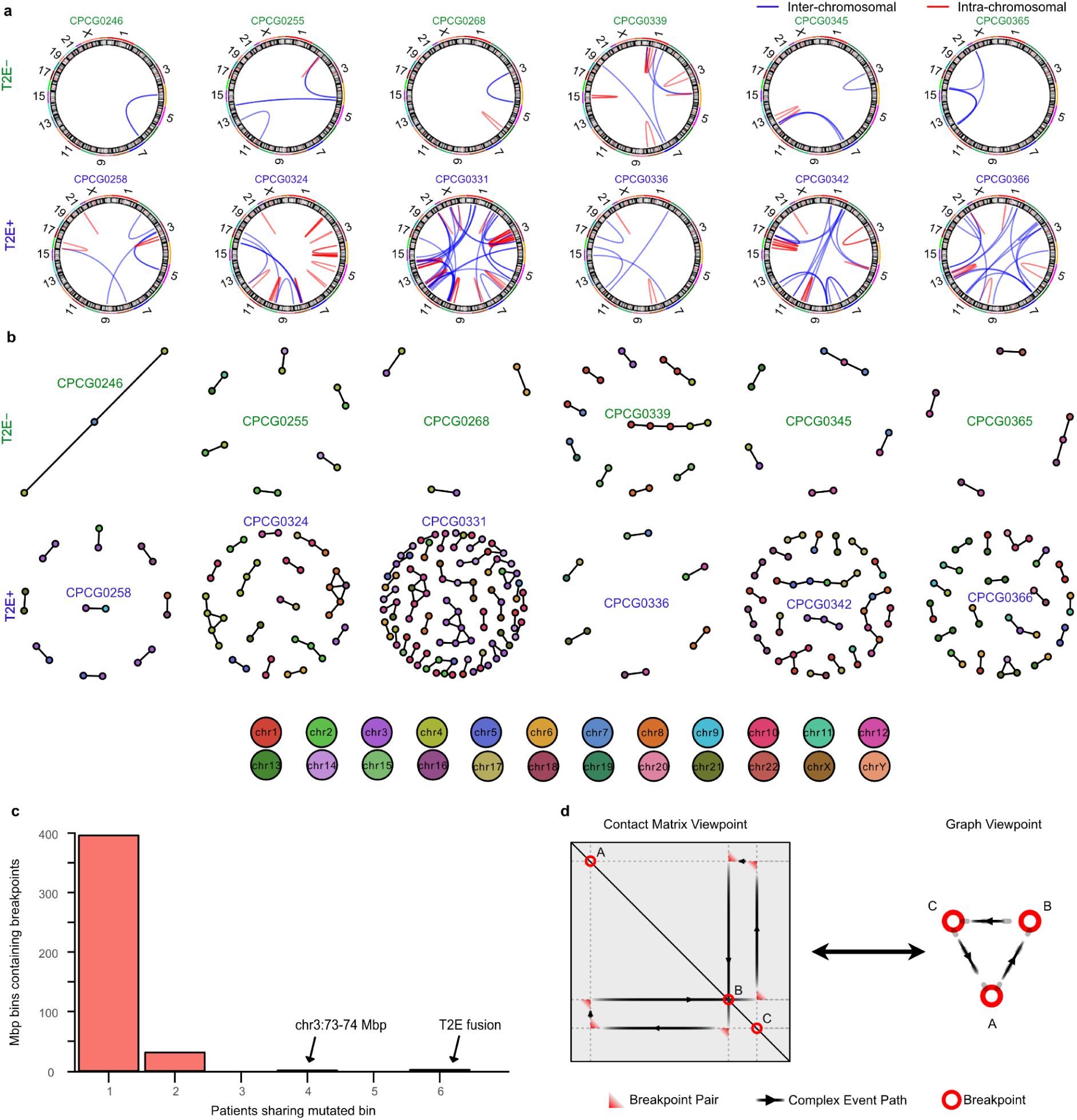
Structural variant detection from Hi-C data. **a.** Circos plots of structural variants identified in the 12 primary prostate tumours. **b.** Graph reconstructions of the simple and complex SVs in all 12 tumours. The node colour corresponds to the chromosome of origin. The nodes are spaced by a spring-force layout which is then adjusted using the Kambda Kawai optimization ^90^ from the NetworkX Python package ^91^. **c.** Barplot of the number of 1 Mbp bins with SV breakpoints from multiple patients. The previously-reported highly-mutated regions on chr3 and T2E fusion are highlighted. **d.** Correspondence between the breakpoint representation in the contact matrices and a graph representation. Each node represents a breakpoint and each edge determines whether the breakpoints were directly in contact, as identified by the Hi-C contact matrix.

**Supplementary Figure 5.**
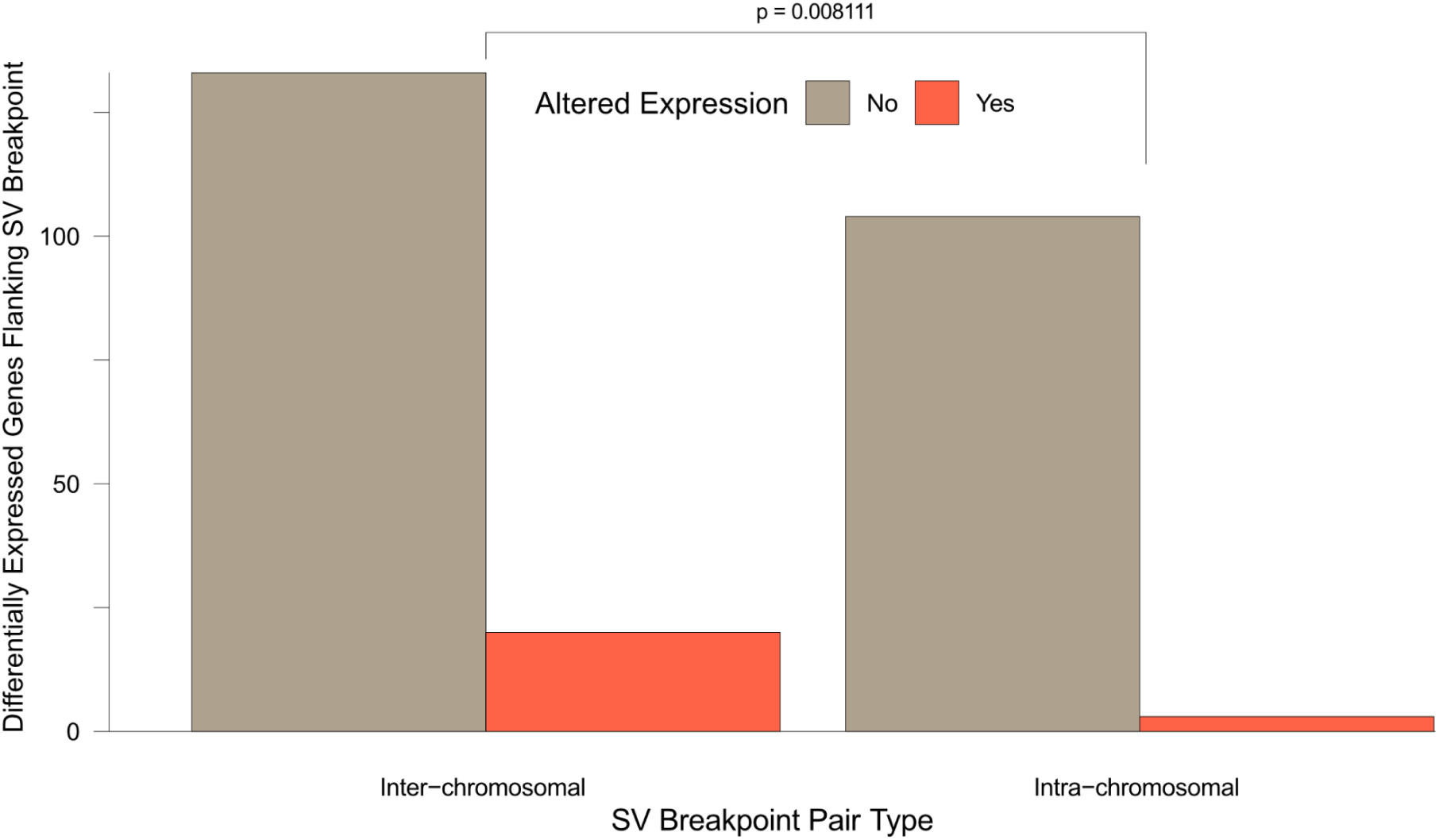
Relationship between inter-chromosomal rearrangements and differential gene expression. Barplot of the number of differentially expressed genes and whether they are involved in SVs spanning multiple chromosomes. Pearson’s chi-squared test, X^2^ = 7.0088, p = 0.00811, df = 1.

**Supplementary Figure 6.**
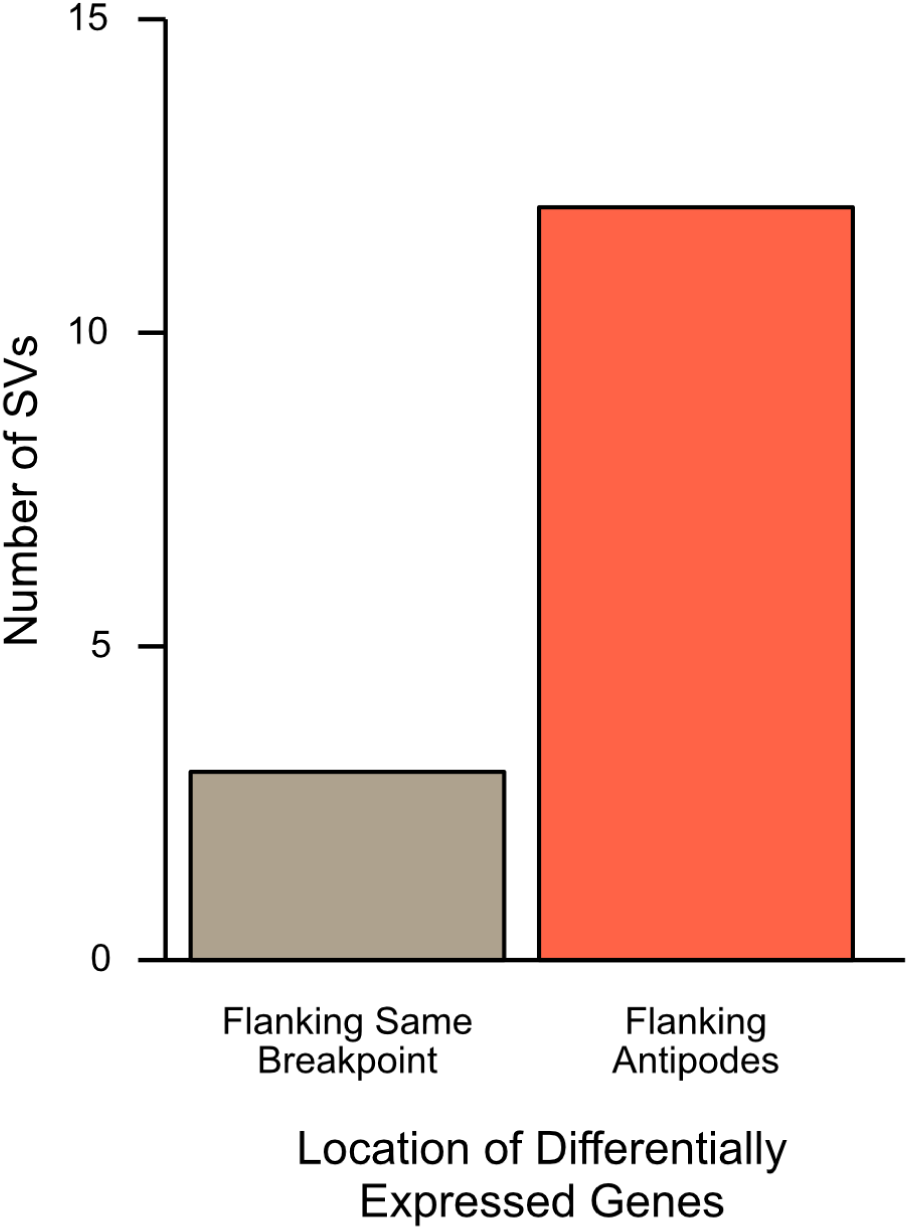
Location of differentially expressed genes around SV breakpoints. Barplot of all 15 SVs associated with both over- and under-expression, categorized by which breakpoints the differentially expressed genes flank.

**Supplementary Figure 7.**
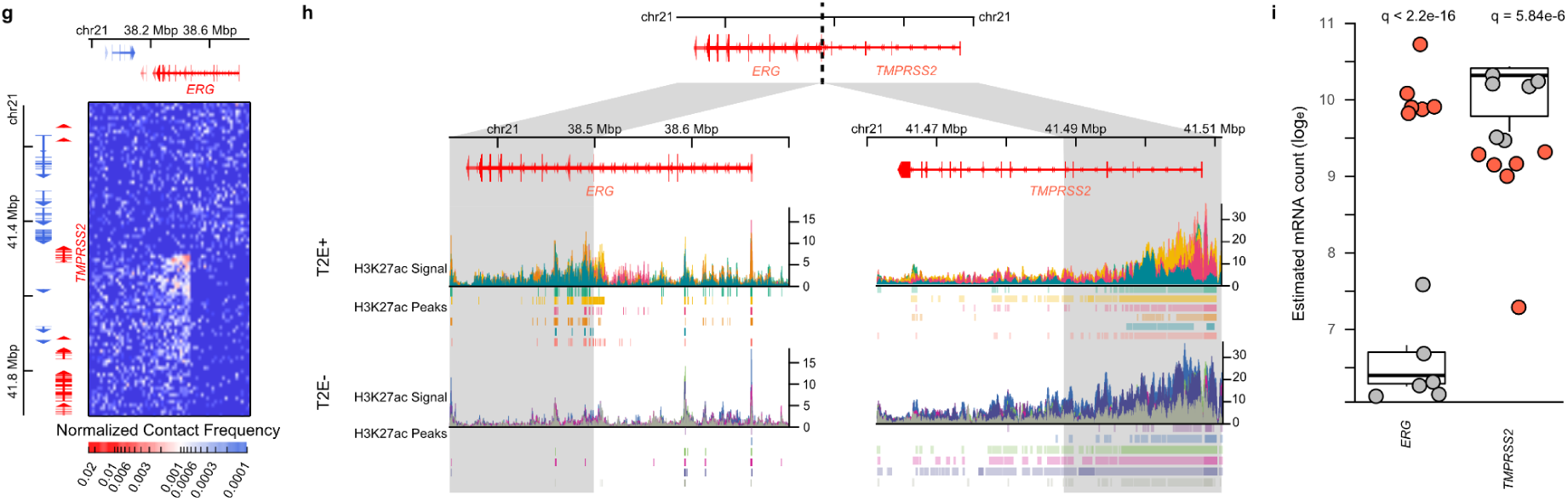
Chromatin organization of the *TMPRSS2-ERG* fusion. **a**. Contact matrix of the deletion between *TMPRSS2* and *ERG.* **b.** Genome tracks of H3K27ac ChIP-seq signal in T2E+ and T2E- patients. The grey region highlights the loci that come into contact as a result of the deletion. **c**. Expression of *TMPRSS2* and *ERG* genes. Boxplots represent first, second, and third expression quartiles of T2E-patients (grey dots). T2E+ patients are represented by red dots.

## Supplementary Tables

**Supplementary Table 1** - Clinical information of samples involved in this study.

**Supplementary Table 2** - Sequencing metrics as calculated by HiCUP for all Hi-C libraries generated in this study.

**Supplementary Table 3** - Summary statistics for TAD counts in all 12 tumour and 5 benign samples, across multiple window sizes.

**Supplementary Table 4** - Individual TAD calls in all 12 tumour and 5 benign samples.

**Supplementary Table 5** - Detected chromatin interactions in all 12 tumour and 5 benign samples.

**Supplementary Table 6** - SV breakpoints detected by Hi-C in each tumour sample.

**Supplementary Table 7** - Simple and complex SVs reconstructed from SV breakpoints.

**Supplementary Table 8** - H3K27ac peaks identified in each of the 12 primary PCa patients. Raw sequencing data as previously published in ^42^ was remapped to the hg38 reference genome.

